# The obesity associated BDNF intron-3 cis-regulatory element, BE5.1, modulates the effects of early-life stress on weight gain, anxiety and gene expression in a sex-dependent manner

**DOI:** 10.64898/2026.09.10.750405

**Authors:** Andrew McEwan, Joseph Morgan, Greg Hutchings, Alexander Rattray, Elaina Collie-Duguid, Jude Al Jishi, Zeynab Heidari, Grace Ritchie, Louie N. van de Lagemaat, Chris Murgatroyd, Alasdair MacKenzie

## Abstract

Early-life stress (ELS) has been associated with increases in obesity and anxiety and affects males and females in different ways. Animal and human studies indicate that ELS has maladaptive effects on brain development due, in part, to disruptions in the expression of genes that include brain-derived neurotrophic factor (*BDNF*). Our previous studies identified a cis-regulatory element (CRE, BE5.1) within BDNF intron-3 that harbours a SNP associated with obesity and anxiety. We used maternal separation (MS) to examine the role of BE5.1 in controlling resilience to the maladaptive effects of ELS. At 9-11 weeks, BE5.1KO mice experience a delay in weight gain compared to WT mice, a phenotype partly reversed by MS in BE5.1KO females. The effects of either MS or BE5.1KO decreased anxiety-like behaviour in males whereas MS had little effect in females who displayed increased anxiety-like behaviour in animals lacking BE5.1. We also found that social behaviours are influenced by BE5.1 deletion in females but are more influenced by MS in males. Furthermore, deletion of BE5.1 and MS together significantly decreased marble burying behaviour in male mice, an effect negated by diazepam. RNA-seq analyses suggest that, in hypothalamus, BE5.1 is required to stabilize or down-regulate the transcription of 26 key genes, with diverse neuroregulatory roles, in response to MS as these 26 genes are strongly upregulated in the absence of BE5.1. We also provide evidence that three of these genes; *Prkcg, Foxg1* and *Mef2c*, modulate BE5.1 activity in an allele-specific manner. Taken together, these results suggest that the BE5.1CRE acts as a buffer element that protects against the maladaptive effects of MS on gene expression and later life behaviours in a sex-dependent manner.

## Introduction

Early-life stress (ELS) is associated with social deprivation and inequality and is known to have significant and lasting effects on physical, mental, and social well-being throughout the life course, being linked to increases in levels of obesity and anxiety in later life [1–3]. Exposure of young people to ELS within a sedentary, obesogenic environment may represent the single greatest threat to health facing young people globally [4]. Furthermore, Western societies are experiencing a major upsurge in cases of anxiety, particularly amongst young people in the wake of the Covid-19 crisis [5]. The economic cost of one form of ELS, referred to as adverse childhood events (ACEs), is estimated at £3 million per individual [6]. If we consider that between 9 and 15% of the UK population suffer four or more ELS events during childhood, this represents a considerable potential burden on Western economies in terms of later-life health effects [7]. In humans, ELS has been associated with internalised disorders such as anxiety and depression in women and externally expressed disorders such as hyperactivity disorder and antisocial behaviour in men [8]. Animal models have also shown that ELS has a significant sex-specific effect on behaviour in later life, such that both males and female C57BL/6 mice demonstrate increased anxiety-like behaviours, and males display hypermobility, increased repetitive behaviours and reductions in social interaction [9]. Understanding the mechanistic links between early-ELS, the later life conditions associated with them, and how genetics, sex and other environmental factors play a role, will greatly facilitate in determining the molecular mechanisms which could be harnessed to support resilience to ELS.

One leading candidate in modulating the effects of ELS is the brain-derived neurotrophic factor (BDNF) protein. BDNF is a common neurotrophin in the central nervous system, responsible for the development and plasticity of neurons by regulating growth and survival, promoting long-term potentiation, and modulating synaptic transmission and activity-dependent plasticity [10]. Many of the effects of ELS are due, in part, to disruptions in *BDNF* gene expression, which plays a vital role in early neural programming and brain health in adulthood [11]. Indeed, various studies have observed a decrease in BDNF expression (mRNA and protein) in several tissues as a result of ELS [12], and ELS is known to alter DNA-methylation of the promoter regions governing BDNF expression in both peripheral human blood and in the tissues of mice [11]. Numerous polymorphisms around the BDNF gene have been associated with obesity [13–16] and anxiety which are conditions exacerbated by ELS [17–19].

Based on these observations, we previously examined the hypothesis that non-coding GWAS-associated SNPs could affect health and disease susceptibility by altering the properties of cis-regulatory elements (CREs) required to control the context-specific expression of BDNF. To explore this hypothesis, we used a combination of comparative genomics and CRISPR genome editing to identify and study polymorphic CREs around the BDNF locus that host disease-associated SNPs [20, 21]. One such candidate CRE lay within BDNF intron 3 (BE5.1), had been conserved for 350 million years and hosted a SNP associated with obesity (rs10767664; p=4.69÷10^-26^; [22]). We found that the A and T variants of BE5.1 drove different levels of activity of BDNF promoter IV (BP4) in primary cells and that analysis of the human UK biobank cohort demonstrated increased association with allelic variants of rs10767664 and anxiety [21]. We reported that deletion of BE5.1 (BE5.1KO) produced a significant increase in amygdala BDNF expression in female mice, who also demonstrated increased levels of anxiety-like behaviour [21]. Further analysis using human eQTL databases (GTex) also highlighted an association with rs10767664 and changes in BDNF antisense gene (BDNF-AS) expression and splicing, and we were able to detect a significant difference in levels of BDNF-AS expression in BE5.1 mice compared to wild type, suggesting that part of the function of BE5.1 was to modulate expression of this gene[21].

In the current study, we explored the possible role of the BE5.1 CRE in modulating later-life responses to environmental challenges such as ELS by maternal separation (MS) and compared weight gain and behaviour of these animals to wild type animals in later life. Significant differences in the effects of MS on anxiety phenotypes, weight gain and social behaviour between wild type animals and BE5.1KO animals were observed and in a highly sex-specific manner. Transcriptomic analysis demonstrated significant changes in the expression of genes in a sex and tissue-specific manner, with particular effects on a group of 26 genes, and was consistent with the hypothesis that BE5.1 acted as a transcriptional “brake” that acted to reduce the deleterious effects of environmental challenge on the nervous system, particularly in the hypothalamus. We further highlight a possible involvement of MEF2c transcription factor binding and protein kinase-C (PKC) pathways in regulating BE5.1 activity in a synergistic and allele-specific manner suggesting a possible mechanism linking the rs10767664 polymorphisms to resilience and disease. We discuss the possible implications of our findings in the context of current understandings of the mechanisms regulating gene environment (GxE) interactions and their effects on human health in later life.

## Materials and Methods

### Derivation and maintenance of animals

BE5.1KO mice were re-derived from frozen embryos and maintained as a homozygous colony on a C57BL/6 background [21]. Animals were maintained under specific pathogen-free (SPF) facilities that conformed to the Federation of European Laboratory Animal Science Associations (FELASA) guidelines, whereby pathogen screening was carried out on a quarterly (3-monthly) basis using sentinel animals. To comply with the latest UK Home Office recommendations and to uphold the principles of the 3Rs (replacement, refinement and reduction), all animals were handled using cupping and tunnel handling techniques instead of the previously used tail handling technique which was shown to induce higher levels of anxiety and stress [23]. In all experiments, male and female homozygous wildtype and BE5.1KO age-matched littermates were housed under standard laboratory conditions (12 h light/12 h dark cycle), in plastic cages with food and water available ad libitum, depending on the experiment.

### Maternal separation

Male and female wild-type mice were paired with other wild-type mice, and male and female BE5.1KO homozygous mice were paired with BE5.1KO mice. Males were removed once clear signs of pregnancy were observed. Once litters were born, a proportion (∼50%) of pups (P0) were removed from their mothers, together with some nesting material, and kept in a heated cage in a separate room for two hours a day for the first two weeks of life.

### Elevated zero maze (EZM)

The EZM consists of an annular dark-gray platform (60 cm in diameter) constructed of opaque Perspex divided into four equal quadrants. Two opposite quadrants were “open”; the remaining two “closed” quadrants were surrounded by 16 cm high dark, opaque black walls. Quadrant lanes were 5 cm in width. Overhead lighting applying 100 lx at the level of the maze. The movement of animals was tracked using a camera and ANY-maze tracking software (Stoelting Europe). Distance travelled, average speed, number of crossings between light/dark zones and freezing episodes, freezing time and freezing latency were recorded.

### Marble burying test (MBT)

The MBT consisted of a “EUROSTANDARD TYPE IV S” (Techiplast) cage (480 x 375 x 210 mm) cage with 5cm depth of wood chip bedding onto which 20 evenly spaced marbles were placed. Animals were then placed in the cage and the numbers of marbles buried after 30 minutes were recorded. This experiment was repeated with groups of animals following i.p. injection of 10 mg/kg of diazepam.

### Three-Chamber Social Interaction Test

This test consisted of a plastic apparatus (60 cm × 40 cm × 20 cm) with an open top divided into 3 chambers (20 cm × 40 cm × 20 cm) by 2 transparent walls and was used to assess social interaction. Each wall contained an open doorway allowing access between the central chamber and the two side chambers. The experimental mouse was habituated to the apparatus for 5 min and a sex matched “stranger” mouse was placed in a cylindrical wired cup (10 cm diameter) in one of the side chambers, alternating per trial to prevent side bias. An identical empty wired cup was placed in an opposite chamber. After the habituation period, the experimental mouse was left to freely explore the apparatus for 10 min. The cumulative time spent in each chamber, as well as the cumulative time spent interacting with an empty wired cup (object) or the wired cup containing the social-partner mouse was measured using a camera and ANY-maze software. Interaction with either was defined as when the experimental mouse was in close proximity (∼ 1 cm) nose-oriented towards the cages[24].

### RNA Isolation and QC Checks

Tissues (amygdala, hypothalamus, hippocampus) were recovered from test animals following euthatol overdose and snap frozen on dry ice for storage. These tissues (10-25 mg) were homogenised in 350 µl Buffer RLT/ DTT solution using needles of increasing gauge (18G, 21G, 23G), whereafter 1 volume of 70% ethanol was added and mixed. Each sample was added to a RNeasy Mini spin column and centrifuged for 15 s at 12,000 *x g*. After which, 700 µl Buffer RW1 was added to each column, centrifuged for 15 s at 12,000 *x g*, whereafter the column was washed twice with 500 µl Buffer RPE, with centrifugation steps of 15 s and 12,000 *x g* performed after each wash. To dry the membrane of the column, an additional centrifugation step of 12,000 *x g* for 1 min was performed. To elute the RNA, 30 µl RNase-free water was added to each column membrane, and centrifuged at 12,000 *x g* for 1 min.

Isolated RNA samples were quantified using Qubit™ RNA HS kit on the Qubit™ fluorometer 3, length analysis were ran using Agilent RNA ScreenTape Assay (5067-5576) on the Agilent TapeStation instrument and purity analysed using NanoDrop™ One Microvolume UV-Vis spectrophotometer.

#### Library Generation using Nanopore cDNA-PCR barcoding kit

To generate each library, Nanopore cDNA-PCR barcoding kit 24 (SQK-PCB114.24) was used. Initially, 10 ng poly(A)+ RNA was added to a 0.2 ml thin-walled PCR tube (made up to 10 µl with nuclease-free water) and mixed with 1 µl cDNA RT adapter and 1 µl annealing buffer, incubated at 60 °C for 5 min, and left to cool for 5 min at room temperature. To each reaction, 3.6 µl NEBNext® quick ligation reaction buffer, 1.4 µl T4 DNA ligase (2M U/ml) and 1 µl RNaseOUT was added, mixed thoroughly. After a 10 min incubation at room temperature, 1 µl lambda exonuclease and 1 µl uracil-specific excision reagent (USER) was added, where each reaction was mixed and incubated at 5 min at 37 °C. Each reaction underwent a clean-up step using RNAClean XP beads, where each reaction was washed twice with 100 µl short fragment buffer and eluted using 12 µl nuclease-free water. To the eluted samples, 1 µl RT primer and 1 µl dNTPs (10 mM) were mixed, incubated for 5 min at room temperature, after which 4.5 µl Maxima H Minus 5X RT buffer, 1 µl RNaseOUT and 2 µl strand switching primer II was added and mixed. After the samples were incubated at 42 °C for 2 min, 1 µl Maxima H Minus Reverse Transcriptase was added, whereafter each sample were incubated in a thermal cycler for 30 min at 42 °C, then at 85 °C for 5 min and consequently cooled to 4 °C.

To 5 µl of reverse transcribed RNA sample, 7.5 µl unique barcode primer (1 µM) and 12.5 µl 2X LongAmp Hot Start Taq Master Mix was added and mixed, where samples were amplified using the conditions found in Table 1.

**Table 1.** Cycling conditions.

| Cycle Step | Temperature (°C) | Time | Cycle No. |
| --- | --- | --- | --- |
| Initial Denaturation | 95 | 30 s | 1 |
| Denaturation | 95 | 15 s | 14 |
| Annealing | 62 | 15 s |  |
| Extension | 65 | 3.5 min |  |
| Final Extension | 65 | 6 min | 1 |

After PCR cycling, 1 µl Thermolabile Exonuclease I was added to each reaction, mixed and then consequently incubated in a thermocycler at 37 °C for 5 min, followed by 80 °C for 2 min. Each reaction again underwent a clean-up step using RNAClean XP beads, where each reaction was washed twice with 100 µl 70% ethanol and eluted using 12 µl elution buffer. Each sample was quantified using Qubit™ 1X dsDNA HS assay kit (Q33230) on the Qubit™ Fluorometer 3 and length and quality checked using Agilent genomic DNA ScreenTape analysis (5067-5365) on the Agilent TapeStation, whereafter samples were pooled together to make 50 fmol equimolar samples in 31 µl (made up using elution buffer). To each pooled library, 1 µl diluted rapid adapter (1.5 rapid adapter stock in 3.5 µl adapter buffer) was added and incubated at 5 min at room temperature.

For sequencing, PromethION DNA flow cells (FLO-PRO114M) were used, where after flow cell priming (flow cell flush/tether), the library solution (32 µl DNA library, 100 µl sequencing buffer and 68 µl library beads) was loaded and run using the super accurate basecalling model, where barcode was trimmed post-analysis.

#### Bioinformatic Workflow

Prior to downstream analysis, quality assessment was performed using Nanoplot (v1.46.1) and FastQC (v0.12.1) to assess the base quality, GC content, sequence length distribution and contamination. After QC, mapping to the mm10 mouse genome (obtained from USCS Genome browser [25] [26] (Kent *et al*., 2002; Navarro Gonzalez *et al*., 2021)) was performed using Minimap2 (v2.30-r1287; [27]) in splice-aware alignment mode (-ax splice -uf -k14) for long read cDNA. For differential expression analysis, gene-level read counts were generated from sorted BAM files using featureCounts (v2.1.1; [28]) using the mm10 RefSeq gene annotation (obtained from UCSC RefSeq [29].

#### Reporter gene Plasmid Design and Assembly

The human BE5.1 region was cloned into the Luciferase reporter vector pTAL-Luc (*Addgene*) using primers designed with 3’ ends complementary to the conserved human BE5.1 sequence, and 5’ ends overlapping the double digested (*SmaI*-HF, *XhoI*-HF (*New England Biolabs*)) p-Luc vector, to create BE5.1(A)pLuc. The BE5.1(T)pLuc and BE5.1ΔMEF2pLuc vectors were designed using additional inside primers to create two fragments in each case (See table 1).

The NEBuilder HiFi (*New England Biolabs: E2623S*) kit was used according to manufacturers’ instructions to assemble the three reporter constructs. 100 ng digested backbone vector was used for each assembly with 1:2 molar ratio vector:insert added to the assembly master mix for each construct. Assemblies were transformed into NEB 5-alpha competent cells and plasmids isolated and purified using GeneJET plasmid midi kit (*Thermofisher: K0481*) according to manufacturers’ instructions.

#### Cell Transfection and PMA treatment

The above plasmids were transfected into the human Neuroblastoma cell line SHSY-5Y. Cultures were maintained and transfected in DMEM/F12 1:1 media supplemented with 10% FBS. Cells transferred to 24 well-plates and transfected at 80% confluence using JetPRIME transfection reagent (*Polyplus: 101000015*) according to manufacturers’ protocol.

Luciferase reporter constructs (BE5.1pLuc vectors) were co-transfected with Mef2C-expressing plasmids (*Addgene: #32515*) to investigate the influence of Mef2c expression on BE5.1-driven Luciferase activity when co-transfected in SHSY-5Y cells. In every instance primary reporter vector and co-expression vector were transfected at a 1:1 molar ratio, with renilla Luciferase-expressing vector (pGL4.74 *-Promega: E6921*) used as a normalisation control at a molar ratio 20:1 (primary vector:renilla). These plasmids were transfected together using jetPRIME reagent at 50 µl jetPRIME buffer, 0.5 µg total DNA, and 1 µl jetPRIME reagent per well (24w plate) at 80% cell confluency.

Phorbyl 12-Myristate 13-Acetate (PMA) was dissolved in Dimethyl Sulphoxide (DMSO) and applied at a working concentration of 50µM. 1 µl was added to each well (containing 0.5 ml growth media) 24 h post-transfection, with 1 µl DMSO added to untreated wells as a vehicle control.

#### Luciferase Activity

Luciferase activity was measured at 48 h post-transfection using the Promega Dual-Glo Luciferase Assay system (*Promega: E2920*) according to manufacturers’ instructions. Growth media was removed and cells washed with PBS before 70 µl Promega Glo Lysis Buffer (*Promega: E2661*) added per well. After 5-10 min 50 µl of cell lysate was moved to a 96w plate (*FisherSci: 10167481*) and 50 µl of prepared Dual-Glo working reagent added per well. 50 µl prepared Stop & Glo working reagent was then added to each well. Luminometer readings were carried out on the Promega GloMax Navigator (*Promega: GM2000*) one minute after addition of Dual Glo reagent and twenty minutes after addition of Stop & Glo reagent as in manufacturers recommended protocol.

#### Statistical Analysis

For animal behaviour and weight gain studies data was statistically analysed using one-way ANOVA or students t-tests using Prism. RNA-seq data was subjected to DESeq2 (v1.50.2) analysis, performed using R version 4.5.2, was used where normalised count data (median-of ratios) with dispersion estimates obtained using empirical Bayes shrinkage. Differential expression analysis was assessed using DESeq2 internal Wald tests, where *p* values were adjusted for multiple testing using the Benjamini-Hochberg procedure to control for the false discovery rate (FDR), hence genes with an absolute log2 fold change (|log2FC| ≥1) with an adjusted *p* value of ≤0.05 were considered statistically significant. Volcano plots were used to visualise differential expression data. To investigate the genotype-dependent transcriptional responses to MS, interaction models were implemented including genotype (WT vs KO), exposure (Control vs ELS) and their interaction term. The model design ∼ genotype + exposure +genotype:exposure or (KO_ELS – KO_Control) – (WT_ELS – WT_Control) was applied. The interaction term asks whether the effect of ELS is different in KO mice models compared to WT mice model, corresponding to a difference-in-differences comparison between conditions. Again, p values were adjusted for multiple testing using the Benjamini-Hochberg procedure (FDR <0.05) and transcripts with an absolute log2 fold change (|log2FC| ≥1) were considered statistically significant.

## Results

In order to determine the role of the BE5.1 CRE in modulating the later life effects of ELS we subjected newborn WT and BE5.1KO pups to maternal separation (MS), a widely accepted model of ELS [30], for two hours a day for the first two weeks of life. After weaning, we studied these mice using several different tests to determine whether the BE5.1 CRE played a role in modulating the phenotypic or behavioural effects of MS in a sex-dependent or independent manner.

### Weight gain and body composition

We first examined the effects of BE5.1 deletion and MS on weight gain and body composition at two time points after weaning (**Fig 1**). MS had no significant effect on weight gain between 7 and 9 of age in male or female WT animals (**Fig 1A and B**), consistent with reports of the “resilience” of C57BL/6 mice in several studies [31–34]. However, deletion of BE5.1 produced a significant reduction in weight gain in both male and female animals (**Fig 1A and B**). In female mice, MS partially reversed the effects of BE5.1 deletion (**Fig 1B**), a phenomenon not observed in male mice (**Fig 1A**). In parallel, we observed a significant reduction in fat mass in male KO animals analysed using echo MRI scanning (**Fig 1C**). This reduction in fat mass, as a result of BE5.1 deletion, was maintained in male animals between 11 to 13 weeks of age (**Fig 1G**) although total weight differences between WT and KO animals failed to reach significance (**Fig 1E**) suggesting a compensatory increase in lean body mass over this time frame. Between 11 to 13 weeks female WT animals demonstrated significant weight gain as a result of MS (**Fig 1F**) although there was a marginally significant reduction in fat mass observed between WT and KO-MS exposed animals (**Fig 1H**).

**Figure 1.**
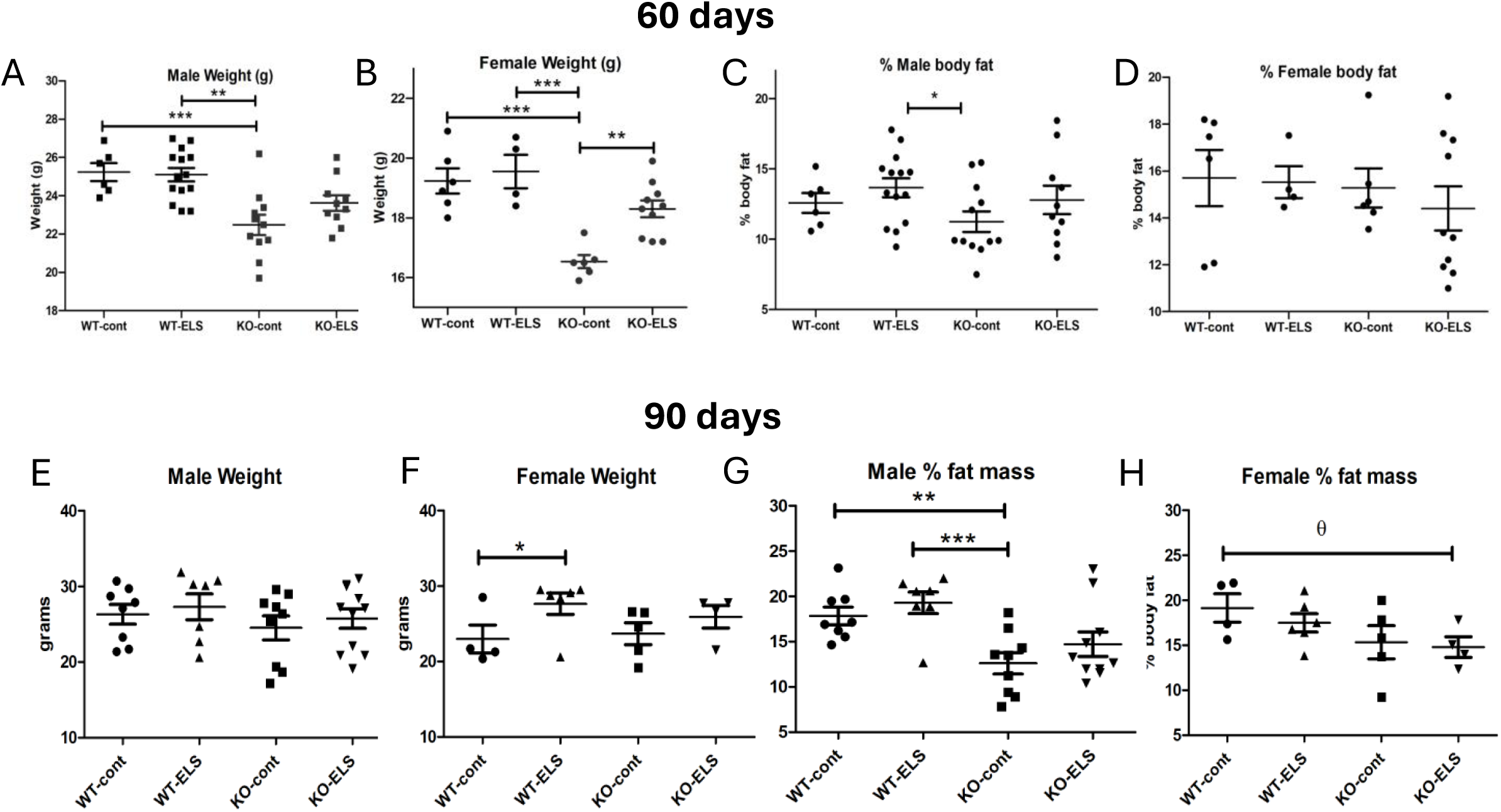
BE5.1 deletion and early life stress have significant effects on weight gain and body composition. (A-H) Weight (g) and body fat composition (% fat assessed using Echo MRI) of male and female wild type (WT) and BE5.1KO (KO) mice that had been raised normally (cont) or following maternal separation (ELS). Weights and % fat mass were assessed on two occasions (60 days old (A-D) and 90 days old (E-H)). Both male and female BE5.1KO mice display a significant delay in body weight increase at 60 days compared to wild types (A and B) suggesting that the BE5.1CRE plays a role in body weight modulation earlier in the life cycle. However, this weight difference is largely negated with age (E and F) although male fat mass remains significantly reduced in KO animals (G). Moreover, ELS results in a significant increase in female weight gain in wild type mice (F) and appears to negate the weight reducing effects of BE5.1 deletion (B). *, p<0.05; **, p<0.001, ***, p<0.005 (one way ANOVA). #, p<0.05, ##, p<0.001 (two tailed t-test). θ, p<0.05 (one tailed t-test).

### Elevated Zero Maze (EZM)

The EZM is designed to explore the balance between an animal’s desire to explore new environments against its anxiety of open space. In male mice, both deletion of BE5.1 and MS increased the time spent in the open quadrants of the EZM (**Fig 2A**) and decreased time within the closed quadrants (**Fig 2B**). The distance travelled overall and in the open quadrants was also increases by both the KO and MS (**Fig 2C and D**). However, there was no significant difference observed in distance travelled in the closed region (**Fig 2E**). The increase in distance travelled in the open quadrants may be partly attributed to the increase in speed displayed by these animals (**Fig 2F**) as well as a significantly decreased immobility time (**Fig 2G**) and distance travelled (**Fig 2C**) within the open quadrants of the EZM. A significant increase in body rotations was also observed in male animals lacking BE5.1 (**Fig 2H**) which suggests a degree of dis-orientation compared to wild type animals.

**Figure 2.**
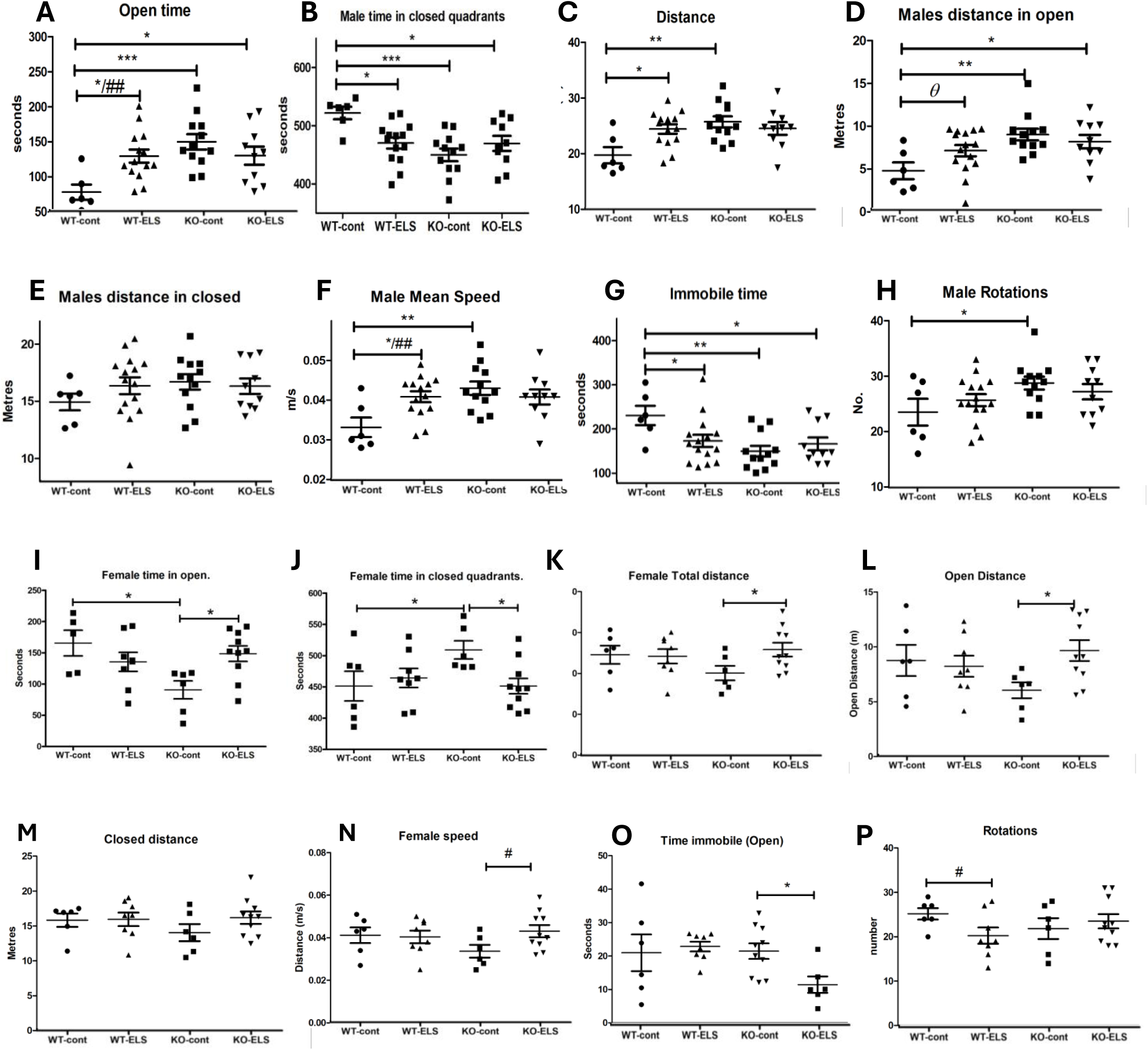
Deletion of BE5.1 (KO) and exposure to maternal separation (ELS) have significant effects on male and female behaviour in the elevated zero maze (EZM). Scatter plots demonstrating the effects of BE5.1 deletion and early life stress on the time (Total test time= 600 seconds) spend by male(A-H) and female (I-P) in the open (**A**) and closed (**B**), quadrants of the EZM. The distance travelled in the open quadrants (**C**), closed quadrants (**D**) and in total (**E**). Male speed overall (**F**), time immobile (**G**) and body rotations (**H**) *, p<0.05; **, p<0.001, ***, p<0.005 (one way ANOVA). #, p<0.05, ##, p<0.001 (two tailed t-test). θ, p<0.05 (one tailed t-test).

Female mice lacking the BE5.1 CRE spent less time in the open quadrants (**Fig 2I**) and more time in the closed quadrants (**Fig 2J**) of the EZM suggesting an increase in anxiety-like behaviour. Deletion of BE5.1 also reduced the total distance covered by females in the EZM in the open quadrants, but this failed to achieve significance (**Fig 2K**). Intriguingly, as observed for weight gain, MS reversed the effects of BE5.1 deletion such that female KO animals exposed to MS demonstrated increased exploratory behaviour in terms of time spent (**Fig 2I**) and distance covered (**Fig 2L**) within the open quadrants of the maze. The increase in distance travelled by female KO mice exposed to MS in open quadrants may be partly explained by the lack of immobility episodes in these quadrants (**Fig 2O**) and the overall increased speed of these animals (**Fig 2N**). MS also decreased immobility time in female KO animals and decreased rotations (**Fig 2O and P**).

Taken together, these results suggest that deletion of BE5.1 has opposing effects in male and female mice such that BE5.1 deletion increases anxiety-like behaviour in females but reduces this behaviour in males who demonstrate increased exploratory behaviour. Intriguingly, MS reinforces the effects of the KO in males and acted to reverse these effects in females in a manner similar to the observed weight loss suggesting the possible triggering of compensatory mechanisms (**Fig 1B**).

### Social interaction test

ELS has been shown to affect levels of socialisation in both human and animal models. To determine the effects of the BE5.1 deletion and MS on social interaction and exploration, we examined the behaviour of BE5.1 and WT mice exposed to MS using the three-chambered social interaction test [24].

Male mice did not demonstrate a significant change in social behaviour as a result of the removal of BE5.1 but social behaviour was significantly influenced by MS **(Fig3A-H)**. For example, MS reduced the distance that males travelled overall (**Fig 3B**) and decreased the number of investigations of the stranger (**Fig 3D**). Male time in the proximal zone and overall % of time in the proximal zone was also affected by MS (**Fig 3E and G**). The distance travelled by WT males exposed to MS was decreased with marginal significance (**Fig 3B**) and head turning was reduced although not to a level that achieved significance.

**Figure 3.**
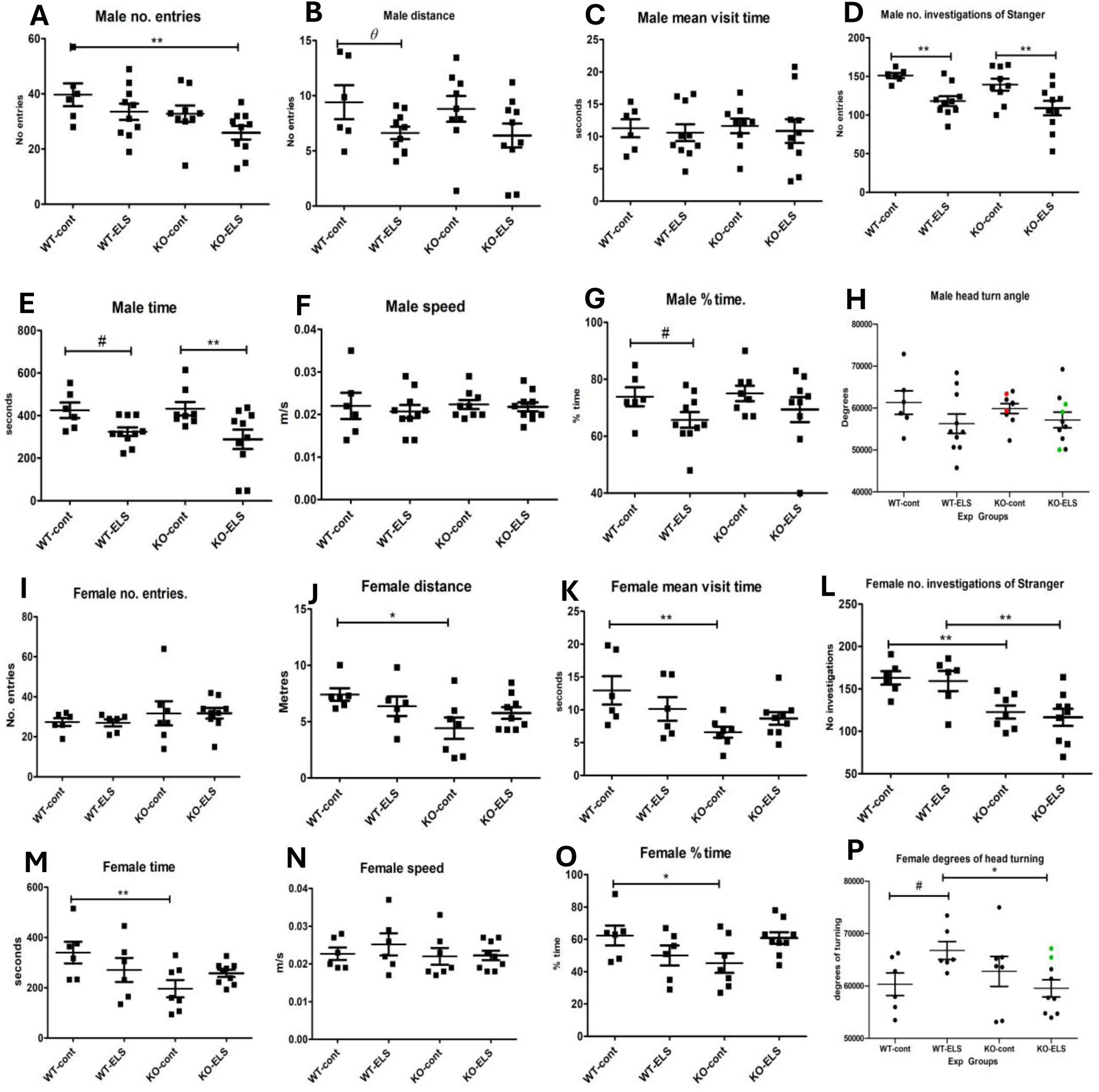
Deletion of BE5.1 and ELS have differential sex-specific effects on social behaviour using the three chambered social interaction test. Scatter plots demonstrating changes in social behaviours in female (A-H) and male (I-P) wild type (WT) or BE5.1 deletion (KO) mice not exposed (cont) or exposed (ELS) to maternal separation. All graphs describe behaviours within a 2cm zone proximal to a sex matched stranger mouse. *, p>0.05, **, p.0.001 by one-way ANOVA. #, p>0.05 by two tailed T-test. θ, p>0.05 by one tailed T-test.

In female mice, MS failed to evoke any significant change in behaviour (**Fig. 3I-O**) except head turning, which was increased significantly (**Fig 3H**). In contrast, significant changes in distance travelled (**Fig 3J**), mean visit time proximal (within 2 cm) of the stranger (**Fig 3K**), number of investigations of the stranger (head directed at stranger whilst in proximal zone), Total time proximal to stranger (**Fig 3E**) and % time proximal to stranger (**Fig 3O**) were all decreased in animals lacking BE5.1 suggesting a role for this element in social curiosity in female mice. Numbers of entries into the proximal zone (**Fig 3I**) and overall speed (**Fig 3N**) were unaffected by either the removal of BE5.1 or MS in female mice.

Taken together, these studies suggest that social interaction behaviours in C57BL/6 mice are modulated by the BE5.1 element in females but not in males. In contrast, male social interaction behaviours are more affected by MS.

### Marble burying test (MBT)

Marble burying is a consistent repetitive behaviour in mice that is often used to model aspects of human conditions with repetitive pathologies such as obsessive-compulsive disorder and autism [35]. Although it has also been used as an anxiety test, its relevance to anxiety is controversial [36]. One of its main values is that it can be used repetitively over a series of several days without a significant alteration, allowing comparisons of different drug treatments. We tested male and female wild-type and BE5.1KO mice that had been weaned normally or subjected to MS with the MBT.

We observed that deletion of BE5.1 and MS had a combined effect in reducing marble burying in males **(Fig 4A)** and decreasing freezing time **(Fig 4B).** Intriguingly, the effects of BE5.1 deletion and MS could be reversed using the anxiolytic positive allosteric modulator of the GABA_A_ receptor, diazepam (**Fig 4C**). A possible role for the GABA receptor is interesting with respect to BDNF modulated behaviours as BDNF is known to modulate the activity of GABAergic system [37] thereby providing a possible clue to the identity of one the systems influenced by MS and BE5.1. Female marble burying was also reduced by the KO (**Fig 4D**), as previously reported [21] but failed to achieve significance. Moreover, MS had no significant effect on marble burying in females (**Fig. 4D**). However, as for males, freezing behaviour was significantly decreased by the KO.

**Figure 4.**
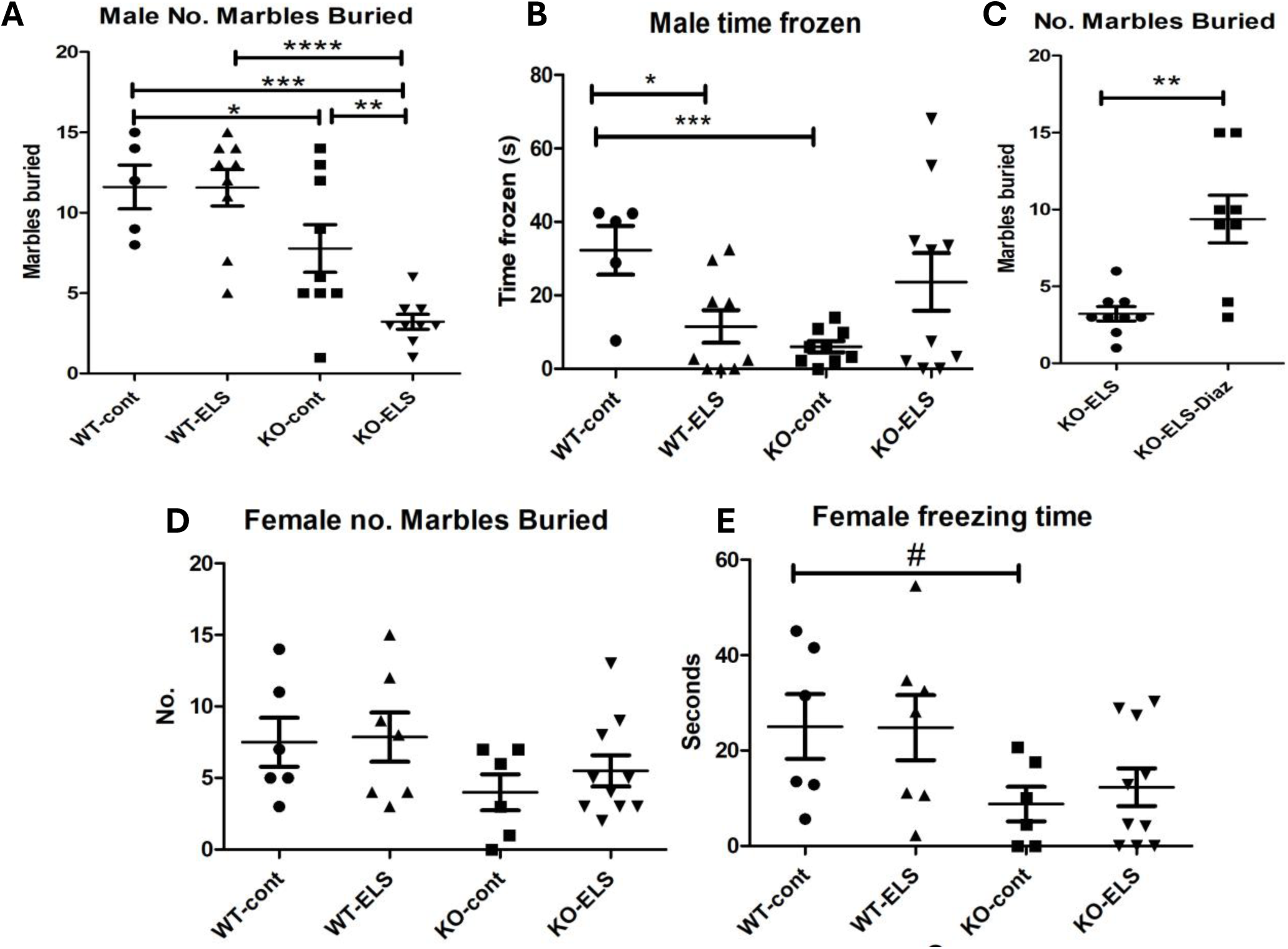
Deletion of BE5.1 and ELS have sex-specific differential effects on marble burying behaviour and responds to GABA_A_ stimulation. Scatter plots demonstrating differences in the numbers of marbles buried by female (A and B) and Male (C-E) Wild type (WT) and BE5.1 knockout (KO) mice as control groups (cont) or following exposure to maternal separation (ELS). E. no marbles buried by male KO animals exposed to ELS and treated with the GABA_A_ agonist diazepam (Diaz). *, p<0.05, **, p<0.001; ***, p<0.005 ****, p>0.0001 by one-way ANOVA. #, p<0.05 by two tailed T-test.

### RNA-seq analysis of gene expression in male and female, WT and BE5.1KO animals exposed to MS

#### BDNF

Based on its location within intron 3 of the BDNF gene we used RNA-seq explore the effects of deleting the BE5.1CRE on the expression of BDNF in WT and KO males and females exposed to MS in 3 different tissues. We observed an increase in BDNF exon 1 expression in pooled WT male RNA (derived from all three tissues) as a result of MS (Fig 5A) which did not occur in KO animals. However, these trends did not achieve significance once adjusted for multiple testing and no other BDNF exon tested showed any change in expression (Fig 5A). We observed a similar increase in BDNF exon 1 expression in pooled tissues in WT females in response to MS and this did achieve significance (Fig 5B). Intriguingly, expression of exon 1 did not change significantly in KO females exposed to MS suggesting a role for BE5.1 in this response (Fig 5B). We also detected a decrease in the expression of BDNF IV in KO females although, this did not achieve statistical significance. Analysis of BDNF expression at a tissue specific level succeeded in detecting a significant up-regulation of BDNF exon1 as a result of MS in male amygdala but this trend was not seen for other exons (Fig. 5C). The expression of exon1 was also decreased in hippocampus and hypothalamus of KO males exposed to MS but this did not gain significance. In female tissues, a similar trend was seen in that MS increased exon 1 mRNA levels in amydala as a result of MS in all three tissues in WT animals but not in KOs. The expression of other exons in female tissues (BDNF exon II and exon IV) also suggested a trend towards decrease expression as a result of BE5.1 deletion although not statistically significant (Fig 5D). Taken together, these data show a trend towards down-regulation of BDNF in the absence of BE5.1 and a reduction in the ability of BDNF to respond to MS.

**Figure 5.**
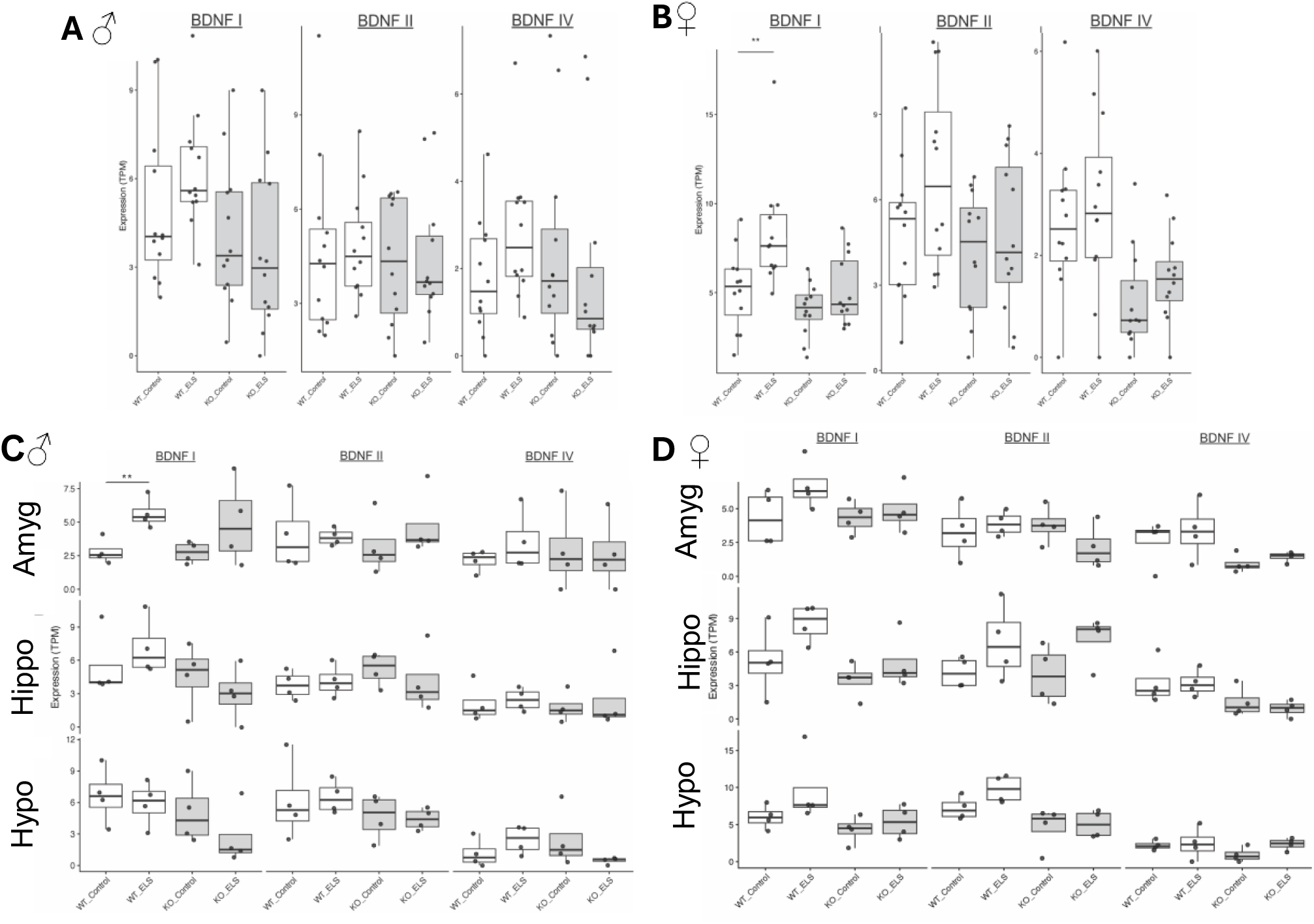
MS has a significant effect on BDNF expression in WT animal but not in BE5.1KO animals. A and B, Sex-stratified BDNF transcript expression derived from RNA-seq data following ELS. Expression levels (measured in transcripts per million (TPM)) of BDNF I, II and IV transcripts are shown across WT and KO models under control and ELS exposure in (A) males and (B) females. Data presented as boxplots with individual points overlaid (pooled brain regions n = 12 per group), where boxes represent interquartile range, with the median indicated by central line. For statistical analysis, Welch’s two-sample (unpaired) t-test was performed. C and D, Brain region-specific BDNF transcript expression in (C) male and (D) female mice following ELS. Expression levels (measured in transcripts per million (TPM)) of BDNF I, II and IV are shown across male (C) and female (D) WT and KO models under control and ELS exposure within amygdala (Amyg), hippocampus (Hipp) and hypothalamus (Hypo). Data presented as boxplots with individual points overlaid (n = 4 per group), where boxes represent interquartile range, with the median indicated by central line. For statistical analysis, Welch’s two-sample (unpaired) t-test was performed. **, p < 0.01.

#### BE5.1 deletion affects the expression of the transcriptome in different tissues and in different sexes at a genome wide level

Given our observations of the effects of BE5.1 deletion on weight gain and behaviours such as socialisation and anxiety, we explored whether BE5.1 might affect the expression of other genes involved in maintaining appropriate brain function in different tissues in male and female animals.

#### Amygdala

The male amygdala only exhibited two significantly downregulated genes in response to BE5.1KO. Enpp2 and Krt18 are both tied to structural or lipid-signalling pathways (**Fig 6A and Supplementary data 1**). By contrast, the female amygdala shows broad, high-magnitude upregulation across hormone signalling (Prl and Gh), mitochondrial metabolism (Mpc1, Hopx, Sgo2a, Gm20594), RNA processing (Rps25, Rps12 Rpph1, Exosc2, Eif3m), and cell-cycle–related genes in response to BE5.1 deletion (**Fig 5B and Supplementary data 1**) suggesting a role for BE5.1 in repressing gene expression. In the absence of BE5.1, the most strongly and consistently upregulated genes include Prl (Prolactin) and Gh (Growth hormone) that are normally expressed in the pituitary. Thus, being expressed in the amygdala suggests altered neuroendocrine signalling, stress-axis dysregulation and changes in hypothalamic– amygdala communication(**Fig 6B and Supplementary data 1**).

**Figure 6.**
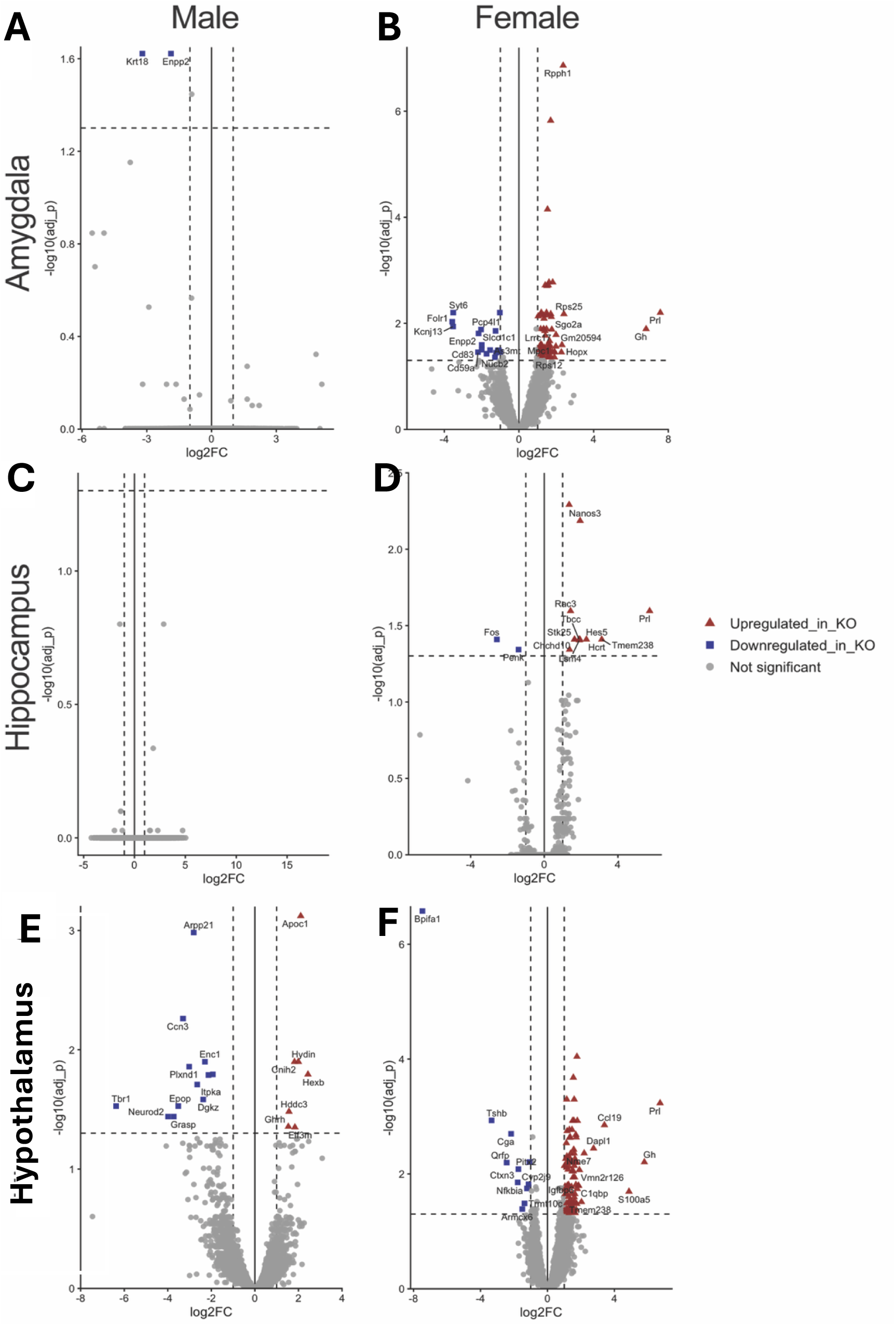
Region-specific BE5.1KO-associated transcriptional differences. Volcano plots showing differential gene expression between KO and WT male and female mice under control conditions across (A and B) amygdala, (C and D) hippocampus and (E and F) hypothalamus brain regions, with log2FC plotted against log10p (Benjamini-Hochberg FDR corrected) values. DESeq2 was ran in R applying a negative binomial generalized linear model with Wald testing (sample size n = 12). Transcripts that reached corrected significance FC thresholds (dotted lines) are highlighted (upregulated FC ≥ 1 = red triangles; downregulated FC ≤1 = blue squares; padj = <0.05). Top 10 differentially expressed genes are labelled.

#### Hippocampus

Whilst no genes in the male hippocampus dataset achieved significance (**Fig 6C and Supplementary data 1**), the knockout of BE5.1 produces a highly sex-biased transcriptional response in the hippocampus, with female mice showing 12 genes up-regulated and two genes down regulated (**Fig 6D and Supplementary data 1**). For example, as seen in the amygdala, the female hippocampus shows robust upregulation of neuroendocrine and neuromodulatory genes in the absence of BE5.1, such as Gh and Prl and in the expression of Hcrt (Hypocretin/Orexin) that is known to regulate wakefulness, arousal and appetite (**Fig 6D and Supplementary data 1**). We also observed an upregulation in the expression of regulators of stem cell maintenance and Notch signalling, Nanos3 and Hes5, hinting at an altered neurogenic or progenitor-like states (**Fig 6D and Supplementary data 1**). This cluster of genes suggests that BE5.1 normally restrains neuroendocrine-related transcription in females. BE5.1KO is also associated with an upregulation of genes involved in cytoskeletal dynamics and cell architecture such as Rac3 ( Rho GTPase controlling actin cytoskeleton and dendritic development), Tln1 (Talin-1, integrin–actin linker essential for adhesion and synaptic structure), Stk25 (encodes a kinase involved in metabolic homeostasis) and Tbcc (tubulin-folding cofactor). With respect to obesity and metabolism Stk25 is interesting as high levels of STK25 expression are associated with increased fatty acid uptake and storage, leading to hepatic steatosis, metabolic dysfunction, and reduced insulin sensitivity [38](**Fig 6D and Supplementary data 1**). Moreover, there is evidence for upregulation of RNA processing and chromatin-remodelling factors including Lsm4 — (mRNA degradation/splicing machinery) and Znhit2 (histone-exchange complex component). Upregulation of Chchd10, a mitochondrial cristae regulator linked to neurodegeneration, may reflect metabolic stress or altered energy demands in the female BE5.1KO hippocampus. Ony two genes are downregulated in female hippocampus further suggesting dysregulation of the neuroendocrine axis. These include Penk (encodes enkephalin, modulates of pain, reward, and stress) and c-Fos (canonical immediate-early gene marking neuronal activation) (**Fig 6D and Supplementary data 1**).

#### Hypothalamus

The effects of BE5.1 deletion on gene expression seems to be most obvious in the hypothalamus of both male and female animals. Male BE5.1KO mice demonstrate a downregulation of neuronal identity transcription factors such as *Neurod2*, *Tbr1*, *Enc1* and *Cplx2* that a are core determinants of excitatory neuron identity. In addition, they experience downregulation of genes encoding signalling enzymes such as *Prkcg Dgkz* and *Itpka* that regulate calcium and lipid signalling, essential for synaptic plasticity and neuropeptide release. This was accompanied by an upregulation of metabolic and ciliary genes such as *Hydin*, *Hexb*, *Hddc3* that mirrors the female metabolic-activation signature (described next), but without the endocrine hormone surge (**Fig 6E and Supplementary data 1**). There is also an increase in the expression of *Ghrh* (Growth hormone-releasing hormone) that is produced in the hypothalamus to stimulate the pituitary gland to produce and release growth hormone (GH) into the bloodstream. Its primary function is to regulate body growth, metabolism, and IGF-1 production.

In the female hypothalamus, we again observed a strong upregulation of pituitary-axis hormones including *Gh, Prl*, as well as *Cga* in the absence of BE5.1 whilst *Tshb* (thyroid-stimulating hormone subunit beta) was down regulated (**Fig 6F and Supplementary data 1**). These changes indicate an alteration in hypothalamic–pituitary signalling, consistent with BE5.1 acting as a regulatory modulator in neuroendocrine neurons. We also see significantly upregulation of mitochondrial genes such as *Atp5md*, *Cox7a1*, *Hint2*, *Mrps6*, *Timm22*, *Tomm40*, *Tkt*, *Mpc1* suggesting increased oxidative metabolism, possibly compensating for altered neuronal firing or peptide release (**Fig 6F and Supplementary data 1**). Intriguingly, there is an increase of Notch pathway components in BE5.1KO females, as well as developmental regulators including *Hes5*, *Crabp2* and *Nkx6-2*. *NKX6-2* is essential for the maturation of oligodendrocytes that produce myelin in the CNS (**Fig 6F and Supplementary data 1**). Mutations in NKX6-2 are associated with hypomyelination and neurological disease [39] in humans. Several neuropeptide gene such as *Qrfp*, *Cga* and *Tshb* are downregulated suggesting that, as these are key hypothalamic peptides controlling appetite, metabolism, and endocrine output, BE5.1 might be normally involved in supporting peptidergic neuron identity in the female hypothalamus (**Fig 6F and C and Supplementary data 1**).

#### Effects of the deletion of BE5.1 on gene expression response to early-life stress

We have demonstrated that deletion of BE5.1 has a significant effect on the expression of many critical genes in multiple regions of the brain that might account for many of the changes in behaviour witnessed in our KO mouse lines (**Fig 1-4**). However, because of the known role of environmental challenge on health and disease risk, we asked whether BE5.1 played a role in modulating the effects of environmental challenge on behaviours that contribute to later disease risk. One such challenge, MS, is a recognised component of social deprivation that has been shown to significantly affect susceptibility to obesity and anxiety in later life in humans. Intriguingly, in behavioural tests (**Fig 1-4**) MS had stronger effects on anxiety and socialisation in male mice than in females who responded more to BE5.1 deletion. To further explore the role of BE5.1 in modulating gene expression in male and female mice in response to MS, and to identify the genetic pathways involved, we explored the effects of ELS by MS on the transcriptomes of hippocampus, hypothalamus and amygdala in male and female mice either weaned normally or subjected to MS.

### Effects of BE5.1KO on MS gene expression response in amygdala

#### Male WT Amygdala

MS elicited a markedly transcriptional responses in the amygdala of WT males with 75 genes being significantly upregulated and 189 being downregulated (**Fig 7A and Supplementary data 2**). The most highly upregulated genes included *Snora30, Mir6236 (both microRNAs), Lars2*, *Bhlhe22* (encodes a transcription factor that regulates neuronal differentiation and maturation) and *Pam* (amidates neuropeptides) but, otherwise, levels of upregulation were modest and included mitochondrial components (*Cpt1a*, *Chchd1*, *Ak3 etc*), non-coding RNA (*Exosc2*, *Snord13*), and neuroendocrine regulators such as *Gh* and *Mas1* (**Fig 7A and Supplementary data 2**). From the perspective of neuro-endocrine signalling, only three genes; *Shisa7* (modulates GABA-A receptor trafficking), *Sema7a* (regulates axon guidance) and *Ttr* (Transports thyroxine and retinol binding protein) were notably downregulated (**Fig 7A and Supplementary data 2**). This pattern suggests a stress-induced metabolic and translational suppression characteristic of a conservative adaptive response. These observations are consistent with reports of the “resilience” against MS previously reported for C57BL6 mice.

**Figure 7.**
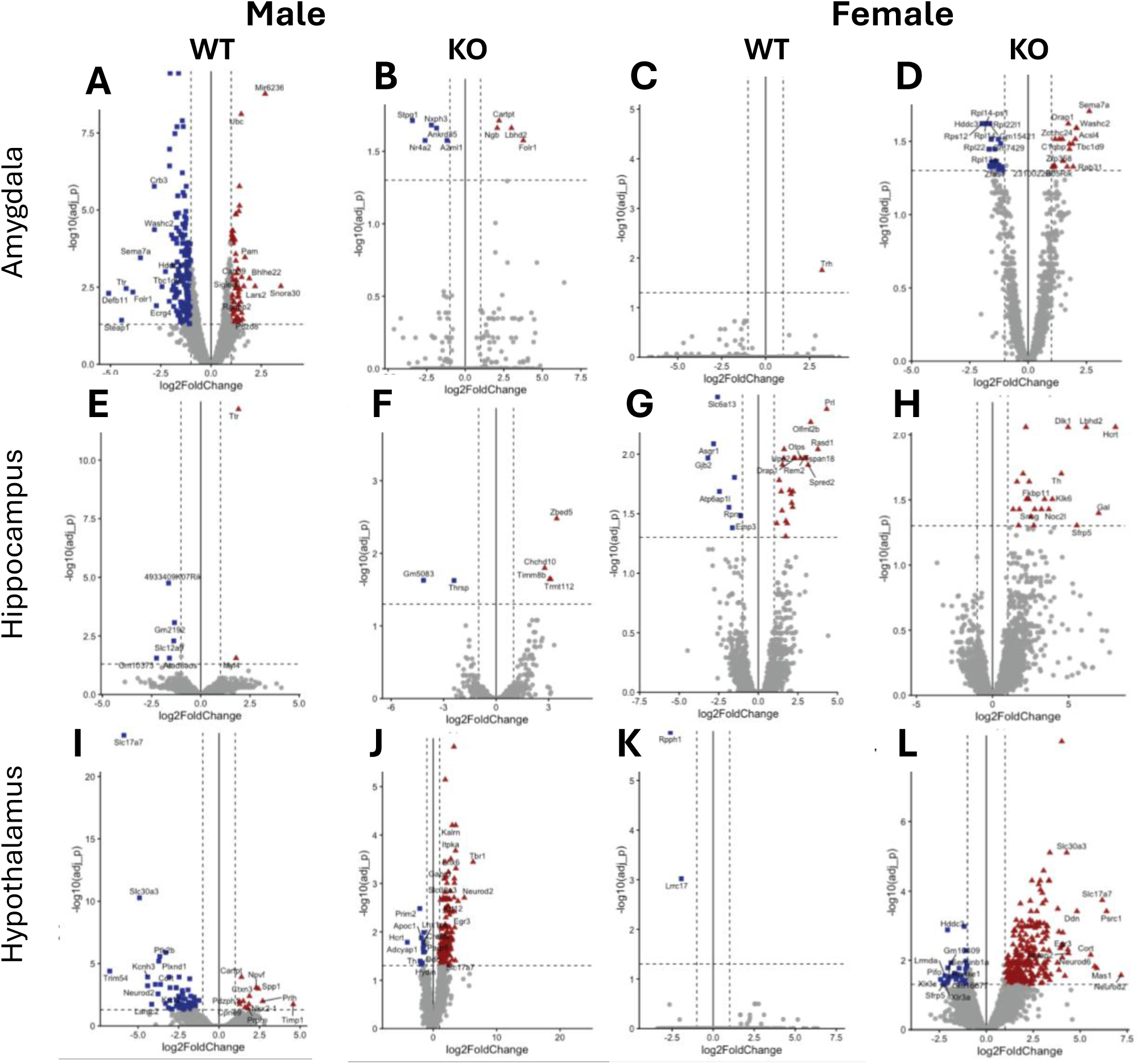
ELS induced changes in gene expression are differentially modulated in WT and KO animals in a tissue and sex specific manner. Volcano plots show differential gene expression between ELS-exposed vs control groups in KO male and female amygdala, hippocampus and hypothalamus, with log2FC plotted against log10p (Benjamini-Hochberg FDR corrected) values. DESeq2 was ran in R applying a negative binomial generalized linear model with Wald testing (sample size per group n = 4). Genes reaching corrected significance FC thresholds (dotted lines) are highlighted (upregulated FC ≥ 1 = red triangles; downregulated FC ≤1 = blue squares; padj = <0.05). Top 10 differentially expressed genes are labelled.

#### Male KO Amygdala

BE5.1KO males did not demonstrate such a broad change in gene expression in amygdala following exposure to MS but the genes highlighted were interesting nevertheless (**Fig 7B and Supplementary data 2**). These include *Cartpt* which was highly upregulated (5.9 log2fold change) and encodes a neuropeptide known to modulate appetite, reward and the stress response. In the amygdala, *Cartpt* is expressed in the CeA and BLA where it strongly co-localises with *PKC*δ, somatostatin and CRF [40]. Another gene, *Nr4a2,* was significantly downregulated and encodes a transcription factor involved in dopaminergic neuron development and survival (**Fig 7B and Supplementary data 2**).

#### Female WT Amygdala

MS produced a minimal transcriptional response in the amygdala of WT females and this response might again reflect the inherent resilience of the C57BL/6 line (**Fig 7C and Supplementary data 2**) with only gene significantly up-regulated; *Trh* (encodes thyrotropin-releasing hormone) whose primary role is endocrine (TSH → thyroid hormones), but also acts as a neuromodulator affecting arousal, stress, feeding, and autonomic circuits (**Fig 7C and Supplementary data 2**). *Trh* is normally highly expressed in hypothalamus (dorsmedial nucleus) and amygdala (Allen Brain Atlas).

#### Female KO Amygdala

KO females exhibited 18 upregulated genes and 30 downregulated genes (**Fig 7D and Supplementary data 2**) and many of the genes in the male WT amygdala, observed to be down regulated following MS (**Fig 7A**, **Table 1 and Supplementary data 2**), are upregulated in the KO female amygdala (**Fig 7D and Supplementary data 2**).

The most interesting genes upregulated in female KO amygdala include *Sema7a*, that regulates axon growth, and *Rtn3,* that plays a role in axonal maintenance (**Fig 7D and Supplementary data 2**). Intriguingly, the vast majority of the 30 genes downregulated in female KO amygdala encode ribosomal subunit proteins suggesting a role for BE5.1 in maintaining protein synthesis in the amygdala following MS. An upregulation in the expression of Trh was not detected in KO animals further suggesting a role for BE5.1 in triggering Trh expression in wild type female amygdala following MS.

### Effects of BE5.1KO on MS gene expression response in hippocampus

#### Male WT Hippocampus

MS elicited a relatively restrained transcriptional response in the hippocampus of WT males, again, possibly reflecting the relative resilience shown by male mice exposed to MS, with only 5 genes down regulated (*Gm2192, SLC12A1, Atad3aos and Gm10373*) and two upregulated (*Ttr and Myl4*) (**Fig 7E and Supplementary data 2**).

#### Male KO hippocampus

In BE5.1KO mice only 4 genes were upregulated (*Zbed5, Trmt112, Timm8b and Chchd10*) and two downregulated (*Thrsp, Gm5083*) (**Fig 7F and Supplementary data 2**). In both WT and BE5.1KO mice, none of these genes appear to play any major role in neuro-endocrine function and suggests that BE5.1 does not contribute to the transcriptional or behavioural changes seen following MS challenge in the male hippocampus. This lack of change in gene expression suggests that the effects of BE5.1 deletion on behaviour is not centred on the male hippocampus.

#### Female WT hippocampus

In comparison, WT females showed an intriguing transcriptional response to MS in the hippocampus compared to males, with neuronally relevant genes such as Spread2 (suppressor of RAS/MAPK signalling), Rem2 (RGK family GTPase), Hpca (Ca^2+^ sensor in synaptic plasticity) Tmem108 (synase organisation) up regulated in expression as well as the previously observed Stk25 and Prl genes further supporting a functional linkage of BE5.1 with these genes. Neuronally relevant down regulated genes included Ptgds (prostaglandin D2 synthase involved in sleep regulation) and Slc6a13 (reuptakes GABA from synapses) (**Fig 7G and Supplementary data 2**). Other neuronally expressed genes that are upregulated in female WT hippocampus in response to MS include Sncg (neurotransmitter release), Scg2 (regulates neuroendocrine signalling), Vat1 (synaptic vesicle function), Gng3 (CGRP signalling) and Podxl2 (synapse formation and adhesion).

#### Female KO hippocampus

Deletion of BE5.1 produced a large increase in the expression of a number of critical neuroendocrine modulators following MS that include Hcrt (orexin), Gal (encodes the galanin neuropeptide involved in feeding and mood behaviour) and Th (tyrosine hydroxylase; a rate limiting enzyme involved in the synthesis of catecholamines (dopamine, noraderenalin etc,) from tyrosine (**Fig 7H and Supplementary data 2**). Unusually, no genes were found to be significantly downregulated in female MS exposed hippocampus lacking BE5.1 suggesting that BE5.1 is required to drive the expression of these genes in the wild-type female hippocampus in response to MS.

### Effects of BE5.1KO on MS gene expression response in hypothalamus

WT males displayed a strong downregulation of 81 genes in gene expression in hypothalamus following MS with only 12 genes being up-regulated(**Fig 7I and Supplementary data 2**). Conversely, only 12 genes were downregulated, as a result of MS, and 167 genes were upregulated in KO males (**Fig 7J and Supplementary data 2**). Only two genes; Nr4a1 and Lrrc17, were significantly affected by MS in WT females both showing only modest downregulation consistent with the previously reported resilience of female C57BL/6 mice to the negative effects of MS (**Fig 7K and Supplementary data 2**). In stark contrast, 284 genes were significantly upregulated in female KO mice subjected to MS, and 34 were down regulated (**Fig 7L and Supplementary data 2**).

Most notably, 26 genes shown to be down regulated by MS in the male WT hypothalamus, were strongly upregulated by MS in KO males and female mice within the hypothalamus **(Fig 8 and 9 and Supplementary data 2**). These included neurodevelopmental transcription factors (*Bcl11b, FoxG1, Neurod2 and Zbtb18),* activity dependent plasticity transcription factors (*Egr1, Egr3 and Mef2c*), synaptic scaffolding and cytoskeleton genes (*Dlgap1, Enc1, Synpo and Wipf3*), Neurotransmitter transporters and vesicles (*Slc7a7, Slc30a3 and Cplx2*), intracellular enzymes and kinases (*Prkcg, Ptk2b, Itpka and Ak5*), Ion channels and receptors such as (*Kcnf1, Kcnh3 and Gabrd*), and neuropeptides and GPCR components (*Cck, Cort, Adcy1, Chrm1 and Camk2a)* **(Fig 8 and 9 and Supplementary data 2**).

**Figure 8.**
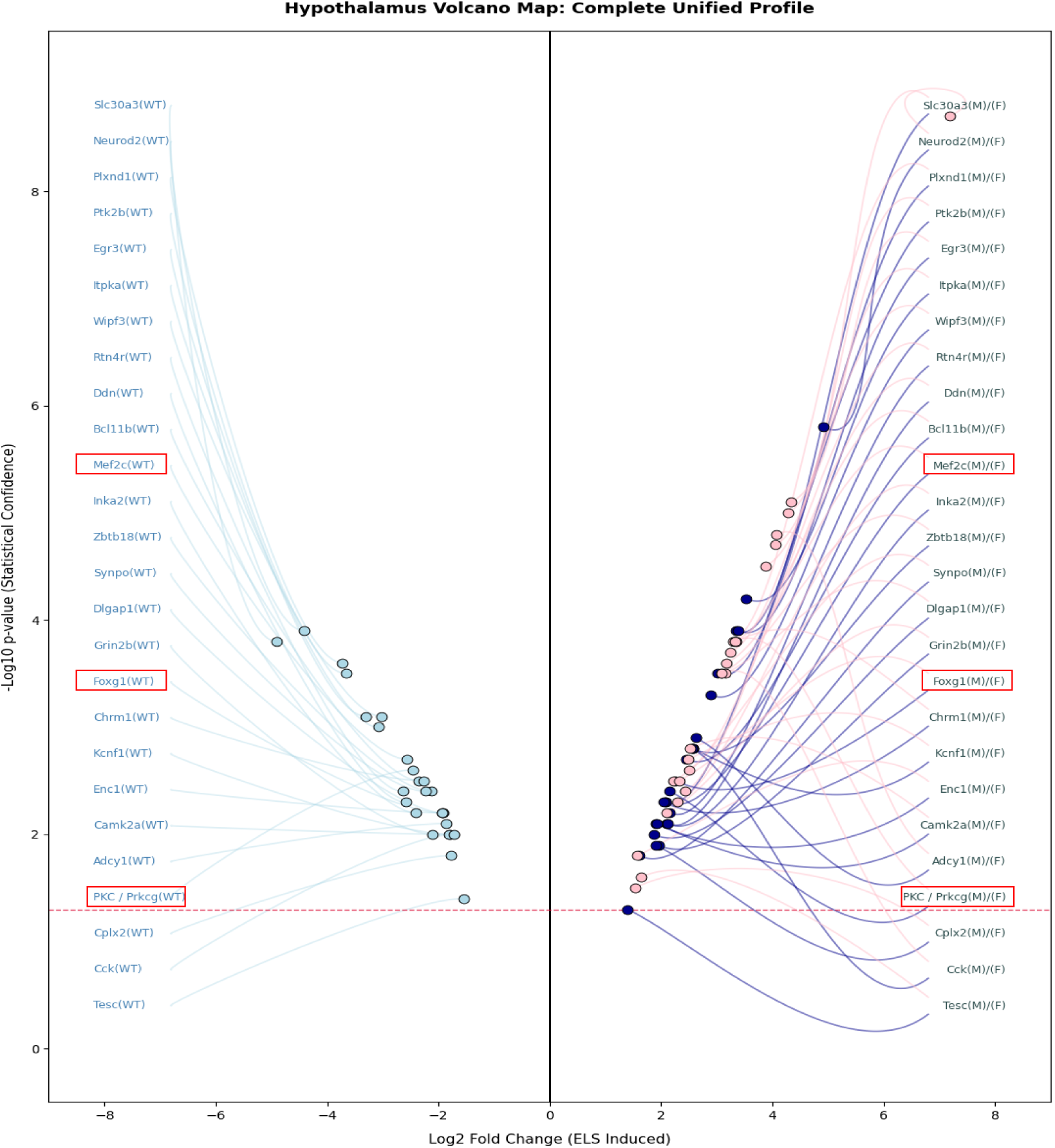
26 genes down regulated in male wild type hypothalamus are strongly up-regulated in male and female KO hypothalamus following ELS. Volcano plots showing differential gene expression between ELS-exposed WT male (light blue), KO-male (dark blue) and KO-female (pink) demonstrating a reversal in the expression of 26 genes in the hypothalamus following maternal separation. Genes relevant to Fig 10 are highlighted (red boxes).

**Figure 9.**
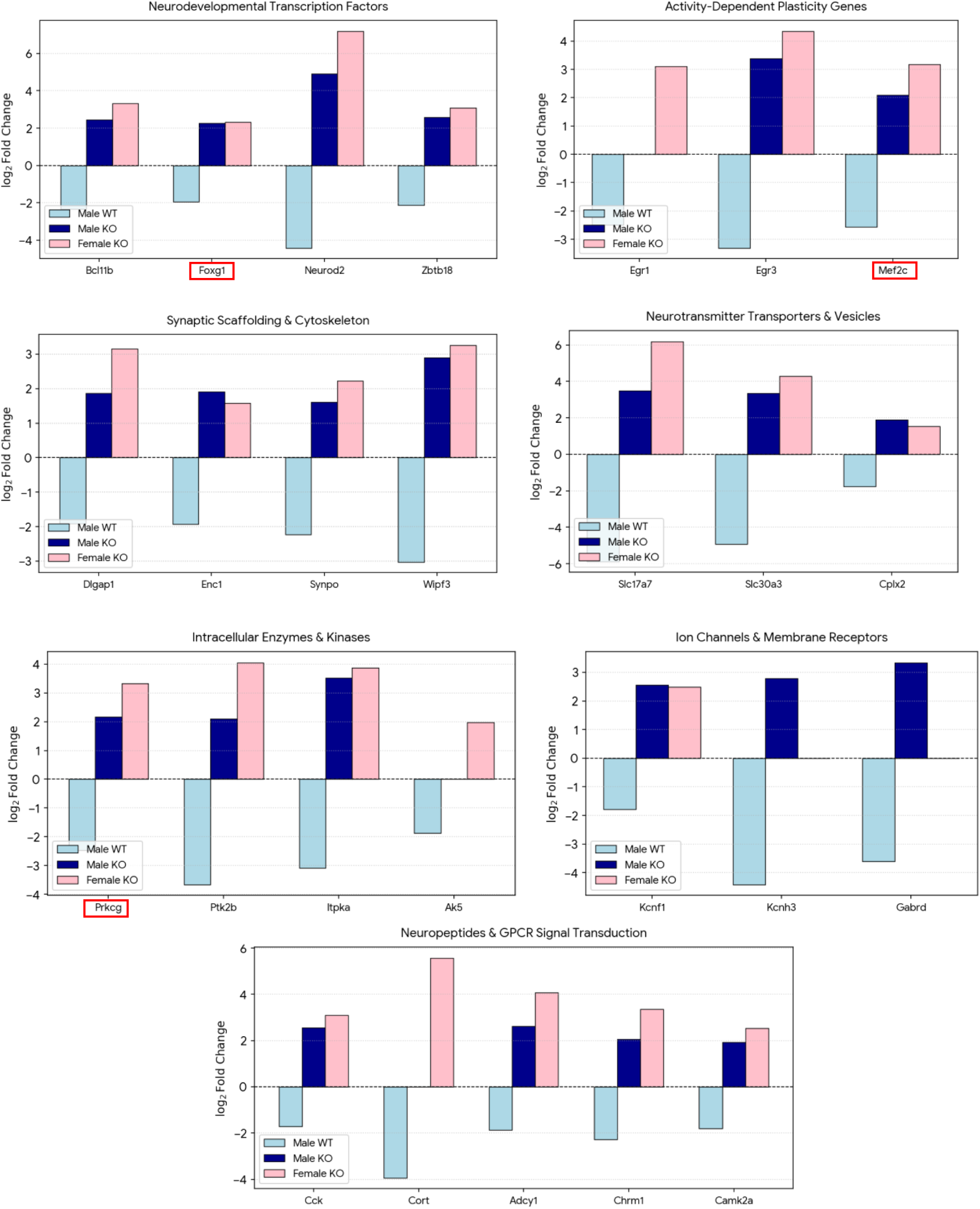
26 genes down regulated in male wild type hypothalamus are up-regulated in male and female KO hypothalamus following ELS. Bar graphs showing differential gene expression, as assessed by RNA-seq and displayed according to function, in the hypothalamus following maternal separation in ELS-exposed WT male (light blue), KO-male (dark blue) and KO-female (pink) mice. Genes relevant to Fig 10 are highlighted (red boxes).

Together, these findings suggest that the BE5.1CRE plays an important regulatory role in shaping hypothalamic sensitivity to early-life stress in WT animals, restraining broad transcriptional activation and maintaining metabolic stability. The expression profiles observed in the absence of BE5.1 suggest the unmasking of a latent stress-reactive network, centred on a discrete group of 26 genes, that were upregulated following MS, fundamentally altering the molecular phenotype of the hypothalamus with an impact on weight gain and behaviour in later life.

#### BE5.1 interacts with MEF2C and responds to Protein Kinase-C (PKC) signalling in an allele specific manner

To gain further clues as to the mechanisms regulating BE5.1 activity, and its role in modulating response to MS, we used the JASPAR CORE 2004 and HMR conserved TFBS tracks of the UCSC browser and identified conserved MEF2 and FOXG1 consensus binding sites within BE5.1 (Fig.12A). This is notable because Mef2c and FoxG1—transcription factors we previously found to be downregulated in the hypothalamus of WT males after MS—are both strongly upregulated in the hypothalamus of both male and female BE5.1KO mice following MS **(Fig 8 and 9 and Supplementary data 2**). Datamining of the UCSC Genome Browser identified two putative MEF2 binding sites (MBS1 and MBS2; **Fig. 10A**) within the BE5.1 enhancer. Notably, MBS2 directly overlaps the rs10767664 (A/T) polymorphism (Fig. **10A and B**), which—uniquely in humans—replaced the T-allele that had been conserved across higher vertebrates for approximately 350 million years (**Fig. 10B**) with the obesity-associated A-allele. We explored the hypothesis that MEF2c interacted with BE5.1 and used site directed mutagenesis to knock out the polymorphic putative MEF2a binding site (MBS2) within a luciferase reporter containing BE5.1 to form ΔMBS2 (**Fig. 10C**)). Co-transfection of a plasmid expressing MEF2c into SHSY-5Y cells with the unaltered BE5.1 increased luciferase expression significantly compared to un-co-transfected cells suggesting a functional interaction between MEF2c and MBS2 (**Fig. 10D**). Moreover, deletion of MBS2 caused a significant drop in BE5.1 activity compared to the wild-type reporter lending support to the hypothesis that both MEF2 binding sites in BE5.1 were functional and that MBS1 acted to partially rescue deletion of MBS2 (**Fig.10D).** RNA-seq analysis of the effects of MS on WT male transcriptome showed a decrease in the expression of the *Prkcg* gene, that encodes the protein-kinase C-protein, in WT male hypothalamus but a greatly increased expression in the hypothalamus of both male and female KOs (**Fig 8 and 9**). This is interesting as *Prkcg* knockout mice display decreased anxiety [41]. We therefore explored the possibility that stimulation of PKC pathways would affect BE5.1 activity in combination with Mef2c expression. Treatment of cells transfected with reporter plasmids containing either the BE5.1(T) or BE5.1(A) alleles with the PKC agonist, phorbol ester (PMA), triggered a small but significant response in reporter expression (**Fig 10E**). In parallel, co-transfection with MEF2c expressing plasmids also upregulated reporter gene expression (**Fig 10E**). However, PMA treatment, when combined with MEF2c expressing plasmid co-transfection, triggered a highly significant increase in reporter BE5.1 reporter gene activity for both the A and T-allelic variants (**Fig 10E**). This observation suggests that the functional interaction of MEF2c with BE5.1 is stimulated by PKC signalling. Interestingly, we observed a significant difference in the degree to which the A and T alleles of BE5.1 responded to MEF2c co-transfection and PMA treatment such that the obesity associated A-allele did not respond as strongly which is consistent with our previous studies in primary hypothalamic cells (**Fig 10E**) [21].

**Figure 10.**
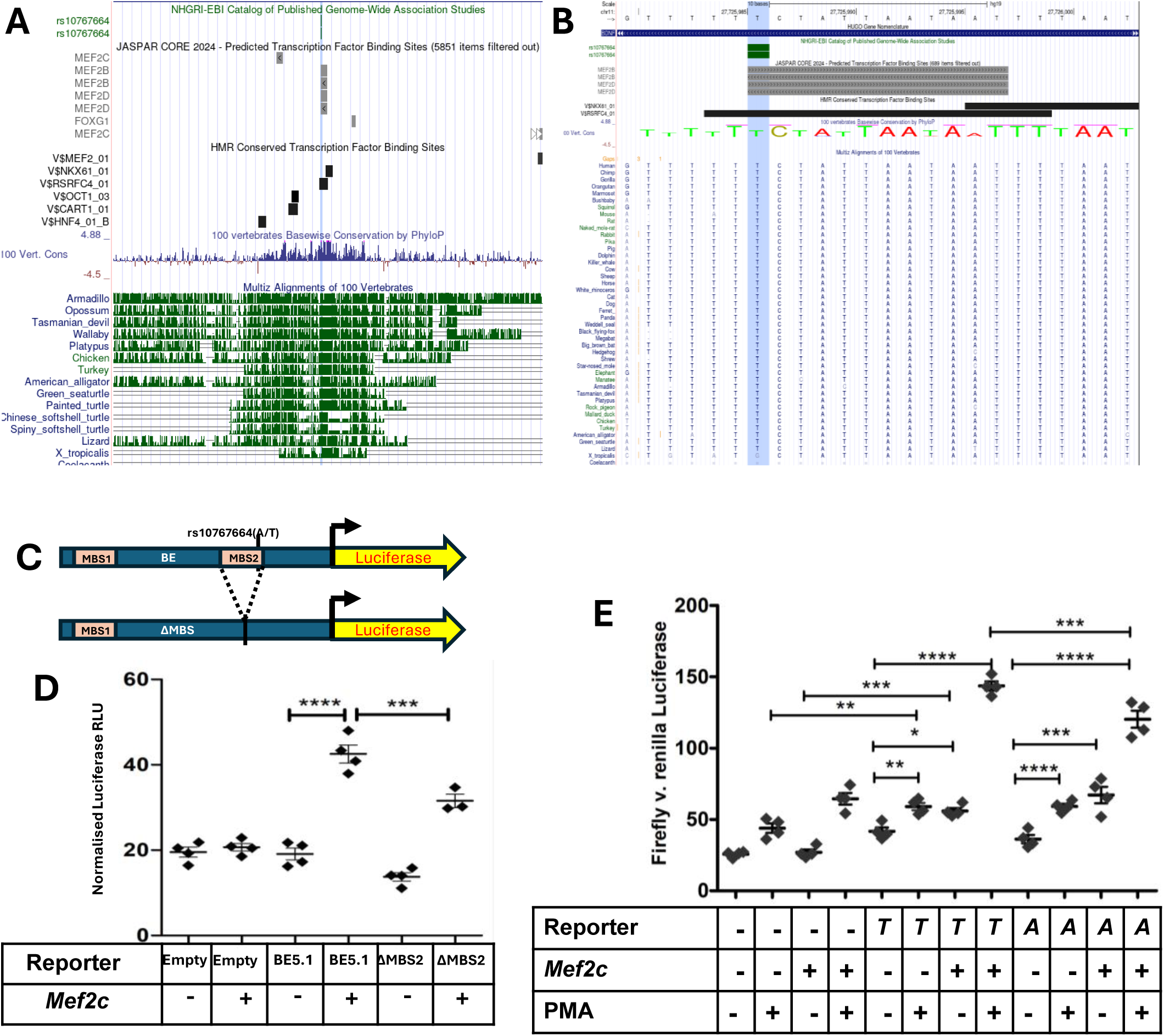
Mef2c interacts with the BE5.1 CRE and interacts synergistically with PKC stimulation to induce allele specific activation. **A.** Screenshot from UCSC browser showing predicted MEF2 binding sites as predicted by JASPAR within the most conserved core region of BE5.1 (Blue peaks and green bars). **B.** Screenshot from UCSC browser showing the relationship of MEF2 binding factor recognition sites (RSRFC4 = Mef2A) at a base pair level and demonstrating the relationship with rs10767664 and multi-species conservation covering 350 million years (perpendicular light blue line). **C.** Diagram showing the reporter plasmids transfected in **D** and **E** (MBS1 and 2; predicted MEF2 binding sites 1 and 2). **D**. Scatterplots showing firefly luciferase expression (normalised to renilla) from empty (empty) reporter plasmids or reporter plasmids containing either wild type BE5.1 or ΔMBS2 transfected into SHSY-5Y cells. Reporters were co-transfected with expression constructs expressing Mef*2c (Mef2c)*. **E.** Scatterplots showing firefly luciferase expression (normalised to renilla) from empty (-) reporter plasmids or reporter plasmids containing either BE5.1T (*T*) or BE5.1A (*A*) allelic variants transfected into SHSY-5Y cells. Reporters were co-transfected with expression constructs expressing Mef*2c* protein. Transfected cells were then treated with phorbol esters (PMA) to stimulate the PKC signal transduction pathway prior to luciferase analysis. *; *p*<0.05, **; *p*<0.01, ***; *p*<0.005; ****; *p*<0.001.

## Discussion

We explored the role of the BE5.1 cis-regulatory element in transmitting the later life effects of ELS by exposing WT and BE5.1 knock out (KO) animals to MS. We found that deletion of BE5.1 delayed weight gain in male and female KO animals at adolescence and that MS partly reversed this in females. We also observed that both the KO and MS reduced anxiety-like behaviour in males but that the KO increased anxiety-like behaviour in females, a response which, as for weight gain, was reversed in KO animals by MS. Deletion of BE5.1 had little effect on socialisation in males which, instead, was affected by MS equally in both KO and WT animals. On the other hand, the KO had a strong effect on the interactions of females to other females, a behaviour unaffected by MS. Males also responded to MS and the KO by burying less marbles; a behaviour reversed by activation of the GABA_A_ receptor. In males, deletion of BE5.1 seemed to predominantly affect gene expression in hypothalamus whereas both the female hypothalamus and amygdala demonstrated strong changes in the expression a large number of genes. We also observed that MS had relatively little impact on gene expression in hippocampus and only had moderate effects in WT males and KO females in amygdala. However, the greatest effects on gene expression were observed in the hypothalamus where a specific set of 26 genes, which were held in stasis or strongly-down regulated in WT animals in response to MS, were strongly upregulated in both male and female animals suggesting that, not only was BE5.1 required to repress the expression of these genes in reaction to stress, but that the majority of its activity was in the hypothalamus. We were also able to provide evidence that three of the genes (Prkcg, Mef2c and Foxg1), out of the 26 observed to be upregulated in KO animals in response to MS, interacted with BE5.1 at a functional level.

### Role for BE5.1 in weight gain

We first explored the effects of deleting BE5.1 on weight gain because of the association of rs10767664 with obesity [22], We previously reported that deletion of the BE5.1 had no significant effect on weight gain between 15 and 30 weeks of age in either male or females [21] and this is consistent with the later time points in the current study. However, examination of weight gain at an earlier time points (7-9 weeks) in males and females suggest that the effects of deleting BE5.1 on weight gain were transient and occurred in during adolescence-a critical developmental period that significantly impacts both the development and the long-term trajectory of weight gain in humans [42]. In males, our RNA-seq analysis suggests that, because deletion of BE5.1 had little effect on gene expression in amygdala and hippocampus, these observed delays in weight gain are likely driven by increased expression of genes such as *ApoC1* and growth hormone releasing hormone (*Ghrh*), that are strongly upregulated in male KO hypothalamus. Moreover, genes involved in brain differentiation, such as *Neurod2* and *Tbr1,* are strongly downregulated in male hypothalamus where they are involved in early brain development suggesting that the observed delay in weight gain may have a deeper developmental origin.

In female mice BE5.1 deletion also caused a delay in weight gain and had relatively little effect on gene expression in hippocampus. However, gene expression in amygdala and hypothalamus responded more strongly to BE5.1KO than in male mice suggesting that the mechanisms affected by BE5.1KO in females was divided between these regions. In females, the most notable changes in gene expression associated with deletion of BE5.1 and delayed weight gain was upregulation of prolactin (*Prl*) Growth hormone (*Gh*) and *Stk25* that occurred in all 3 tissues, and strong down regulation of *Qrfp* and Thyroid-stimulating hormone beta subunit (*Tshb*). Changes in the expression of these genes could account for the difference in weight gain observed between female WT and KO animals at 7-9 weeks.

### Role for BE5.1 in anxiety

Deletion of BE5.1 decreased anxiety-like behaviour in males (increased time in open quadrants). This observation could be skewed by the observation that KO males travelled a greater distance, faster and with less immobile episodes whereas female speed and distance were not as strongly affected. Gene expression in male amygdala and hippocampus did not change significantly in response to BE5.1KO with only the hypothalamus demonstrating any significant change. Thus, the decrease in male anxiety-like behaviour or the increase in movement and exploratory behaviour observed is likely to be centred on the hypothalamus and could be function of increased expression of genes including *Hexb*, and decreases in expression of *Prkcg*(Protein kinase C gamma) [41], Dgkz (Diacylglycerol kinase zeta) that modulates PKC pathways[43], or *Arpp21* [44] that have a demonstrated role in anxiety/stress modulation.

A more dynamic response to BE5.1 deletion was observed in the female brain where gene expression was strongly affected in all tissues examined. Thus, the increased anxiety-like behaviour displayed by females was associated with ↑*Prl,* ↑*Gh,* ↑*Cck* ↑*Cort* and ↓*Nucb2* and ↓*Enpp2* in amygdala ↑*Prl,* ↑*Hcrt* and ↓*Penk* in hippocampus and ↑*Prl,* ↑*Gh,* ↓*Qrfp* and ↓*Tshb* in hypothalamus many of which are associated with anxiety phenotypes [45–47]. Human [48] and animal studies [49] have demonstrated a role for elevated prolactin in increasing anxiety. It is also possible that the exogenous mis-expression of *Prl*, in combination with a number of other genes due to BE5.1 deletion, may have tipped the balance of homeostasis in favour of increased anxiety-like behaviour observed in KO females in the EZM. Moreover, increased expression of orexin has been linked to modulation of the stress response and anxiety[50]. Thus, mis-expression of the *Hcrt* gene, as a result of BE5.1KO, in all female tissues examined, could also contribute to the increased anxiety response observed. However, because all these tissues form a major part of the limbic system that controls mood, it would be hard to conclude, with any certainty from the current data, which tissues changes in gene expression, made the greatest contribution to the observed mood phenotype.

We previously reported that deletion of BE5.1 increased anxiety-like behaviour in female mice but had comparatively little effect on male mice using the elevated zero maze [21]. The current study confirms these previous studies in females but differs in male mice such that male KO mice demonstrate significantly reduced levels of anxiety-like behaviour compared to WT. Since our previous study [21], our animal facility has adopted a revised animal handling procedure, referred to as “cupping and tubing”, that avoids tail handling and espouses moving animals using cupped hands or opaque plastic tubes which greatly reduce stress in these animals [23]. Thus, we cannot exclude the possibility that this revised handling technique might have reduced overall generational stress on these animals thereby revealing a greater contrast in behaviour between male WT and KO animals previously masked by generational stress caused by tail handling. This hypothesis is supported by the clear evidence of the stress sensitivity of the BDNF gene in the presence of BE5.1 as well as the effects of MS on behaviour and gene expression.

### Social and repetitive behaviours

Deletion of BE5.1 had little discernible effect on male social behaviour consistent with the relatively weak changes in gene expression observed in all tissues. However, deletion of BE5.1 had a much stronger effect on female interaction with sex matched stranger animals such that the distance travelled next to the stranger, as well as the time spent exploring the stranger zone, were reduced in the absence of BE5.1. This suggests that social behaviour is more genetically “hard wired” into the brains of female mice and that male behaviour is more influenced by environmental challenges. Both males and females also displayed a slight reduction in interest in marble burying although without significance. Intriguingly, both sexes displayed a significant reduction in freezing that may reflect a reduction of Prkcg (encodes PKC) expression in male hypothalamus, known to be associated with reduced freezing behaviour [51].

### Weight gain

Sex differences in weight gain as a result of early-life stress have been recognised in both humans and rodent models [52]. Although, the effects of maternal separation (MS) on behaviour, as a model for early-life stress, vary considerably between mouse lines, the consensus is that the C57BL/6 mouse strain is relatively resilient to the effects of MS [31–34, 53, 54]. Our current studies are consistent with these observations showing little or no change in the weight of wild type males or females as a result of MS [21]. Intriguingly, however, we identified a significant reversal in the effects of the BE5.1KO in females exposed to MS suggesting the triggering or de-repression of a pathway normally repressed by BE5.1 activity. This de-repression could be associated with the upregulation of genes that include *Cck* that directly modulates satiety [55] as well as *Mef2c* and *Prkcg* in the hypothalamus, that are known to be involved in the development of reward pathways. Although deletion of BE5.1 significantly delayed weight gain in male mice, MS failed to significantly reverse the effect of the KO seen in females. This suggests that, despite the major upregulation in gene expression observed in male MS exposed KO hypothalamus, redundancies and contingency pathways act to stabilise weight gain in the face of environmental challenge in these individuals.

### Anxiety-like behaviours

Although previous effects of MS on anxiety-like behaviour in C57BL/6 mice suggest resilience in this strain [54, 56, 57] more recent studies report decreased anxiety following MS exposure [34, 58]. In line with these latter studies, we observed decreased anxiety in WT males previously subjected to MS which was accompanied by a large change in the expression of 263 genes in the amygdala consistent with the phenotypic changes observed. In contrast, MS altered the expression of relatively few genes (10) in KO animals, consistent with the relatively small difference on the effects of MS on anxiety-like behaviour in these animals. These findings suggest that BE5.1 contributes to the normal male amygdala response to maternal separation, by altering the expression of 263 genes, and that deleting BE5.1 markedly blunts this response in KO males — both in anxiety-like behaviour and in the underlying gene-expression changes.

WT females showed no significant change in anxiety-like behaviour after MS, consistent with the absence of MS-induced gene-expression changes in amygdala. In contrast, female KO mice exposed to MS showed a significant decrease in anxiety-like behaviour that coincided with a strong upregulation of 18 genes and robust downregulation of 30 predominantly ribosomal genes. This intriguing observation is consistent with the known effects of stress induced glucocorticoids that downregulate ribosome production via the mTOR signalling pathway [59]. Thus, BE5.1 may provide a protective function that helps maintain translational capacity in the amygdala following MS. Interestingly, 10 of the 18 genes upregulated in MS exposed female amygdala, were strongly down regulated in WT males exposed to MS suggesting that BE5.1 holds a central position in the regulation of these genes in the amygdala in response to MS.

Neither WT or KO males demonstrated much of a change in gene expression in hippocampus following MS suggesting that BE5.1 does not affect stress modulated behaviour via the hippocampus in males. In comparison, WT females showed a relatively strong transcriptional response to MS that, again, involved upregulation of Prl and Stk25 and down regulation of prostaglandin synthase (Ptgds) and GABA transport (Slc6a13). In addition, MS triggered high levels of expression of hypocretin (HCRT) and Galanin (Gal) in female KO hippocampus both of which are involved in feeding behaviour. Thus, up regulation of these neuropeptide genes in the hippocampus may contribute the reversal in weight gain seen in KO females exposed to MS.

One of the most striking observations to come out of the current study was that a set of 26 genes, that were either downregulated (male) or held in stasis (females) by BE5.1 in wild type animals in response to MS, but were strongly upregulated within the hypothalamus of KO males and females following MS. Because the hypothalamus regulates autonomic function, metabolism, and the neuroendocrine response to stress, the transcriptional shifts observed following MS may have several biological ramifications. For example, *Slc17a7* (VGLUT1) and *Slc30a3* (ZnT3) undergo a transcriptional swing from baseline suppression in WT males and females to high upregulation in the absence of BE5.1. VGLUT1 controls the packing of glutamate into presynaptic vesicles [60], while ZnT3 packages zinc, a critical co-released modulator of post-synaptic NMDA receptors [61]. The switch from down regulation to upregulation also suggests a change in neurotransmitter dynamics. It is possible that the tissue moves from a state of restricted glutamate release to a hyper-glutamatergic state characterized by significantly higher vesicle loading and release capacity that, in the absence of BE5.1, could cause localized hyperexcitability within hypothalamic nuclei, lowering the threshold required to activate downstream neural circuits.

Both *Prkcg* (PKCγ) and *Ptk2b* (Pyk2) are upregulated following MS in male and female KO in the hypothalamus. Pyk2 is a non-receptor tyrosine kinase activated by calcium influx, which downstream potentiates NMDA receptor channels [62, 63] while PKCγ acts as a primary driver of AMPA receptor trafficking and its expression increases anxiety [41]. Thus, upregulation in the expression of these two enzymes might prime the post-synaptic machinery. Even minor presynaptic inputs may trigger amplified cellular cascades, reinforcing local synaptic transmission and long-term potentiation (LTP). This could represent a transition from a post-synaptic signalling deficit, that contributes to resilience to environmental challenge, to a state of heightened cellular sensitivity with behavioural consequences.

Expression of scaffolding genes like *Dlgap1* (GKAP), *Synpo* (Synatopodin), and *Wipf3* is also repressed by BE5.1 following MS. Synatopodin is essential for the structural integrity of the dendritic spine apparatus [64], while GKAP links post-synaptic receptors directly to the underlying actin cytoskeleton and has been associated with anxiety-like and obsessive-compulsive behaviours [65, 66]. Thus, the up regulation of these structural elements implies active, physical remodelling of synaptic connections. Dendritic spines may undergo significant morphological changes, shifting toward denser or more stable micro-anatomical networks. This points to a structural rearrangement of hypothalamic wiring that may work in concert with the shift in chemical signalling to alter behaviours.

Core endocrine signalling elements like *Adcy1* (Adenylate Cyclase 1), *Chrm1* (Muscarinic Acetylcholine Receptor M1), and neuropeptides like *Cck* and *Cort* shift entirely into positive expression. Thus, because Adenylate Cyclase 1 couples receptor activation to cAMP generation, its upregulation could conceivably provide a boost to secondary messenger signalling. This may hyper-sensitize neuroendocrine pathways, which can disrupt homeostatic feedback loops and alter the baseline set points of downstream autonomic and stress-response circuits [67].

### Social interaction, repetitive behaviour and hyperactivity

The effects of BE5.1 deletion and MS on sociability, as determined using the three-chambered social interaction test, also varied considerably within the literature, with studies in C57BL/6 mice showing reduced social interaction in females and resilience (no change) in males [68]. However, more recent studies recognised a significant reduction in social interactions in males exposed to MS [69]. In the current study we observed resilience in females (little or no change) following MS but a significant reduction in social behaviour in males. In contrast, deletion of BE5.1 induced a significant reduction in female social behaviour whilst having little effect on the interaction of male mice with stranger mice. This clear sex-specific difference suggests that social behaviour in female mice is genetically hard-wired and depends on molecular mechanisms of which BE5.1 is a key component. The importance of BE5.1 in female behaviour was previously reported [21] and parallels its role in anxiety-like behaviour as determined using the EZM suggesting that, in female mice, there is a strong overlap in the BE5.1-dependent mechanisms controlling anxiety-like and social behaviour. In males, however, our observations suggest that anxiety (BE5.1 dependent), and socialisation (BE5.1independent) are modulated by pathways that may not overlap to the same degree as that seen in females. Similar inconsistencies exist for exploratory behaviour, where MS is shown to decrease or increase exploratory behaviours depending on the specific test apparatus [70].

A link between early-life stress and conditions such as obsessive-compulsive disorder (OCD) [71] or autistic spectrum disorder (ASD) [72] has been well established, and these disorders are often co-morbid with anxiety. The role of BDNF in this interaction has been suggested based on a number of different studies in humans [73, 74] and mouse models [75]. The Marble burying test (MBT) is a widely used method to assess repetitive behaviours reflective of aspects of OCD or ASD behaviours [76] although its role in assessing anxiety states is questionable [36]. With regard to the effects of MS on MBT in mice, although there is considerable variation between different strains of mice, it is generally held that C57BL/6 mice are resilient to the effects of MS on marble burying behaviour as seen in other strains such as BALBc [77] [76]. Our observations in the current study are consistent with these studies in that we see no effects of MS on the burying of marbles by either wild type C57BL/6 males or females. Although we observed an insignificant reduction in the numbers of marbles buried in the MBT by males and females lacking BE5.1 this reduction achieved strong significance when combined with MS in males. This intriguing observation suggests that BE5.1 is an important component in the regulatory systems which drive repetitive marble burying behaviours in male mice and, again, implies that BE5.1 acts as a buffer against the effects of MS. Another important property of the MBT is that it is that, whilst animals become habituated to the EZM, the MBT can be carried out multiple times on the same mouse to yield similar results. In addition, MBT activity can altered using drugs targeting the GABAergic or serotonergic systems in mice [78]. This reversal in behaviour indicates that components of the mechanisms that involves BE5.1 in repetitive behaviour might involve the GABA_A_-receptor and provides us with clues as to their identities.

### Evolutionary advantages of conserving the BE5.1 element?

The BE5.1CRE element has been highly conserved over 350 million years of divergent evolution, and one has to ask why. One evolutionary advantage of conserving BE5.1 centres on its function as a conditional genetic buffer or transcriptional "brake". In the wild, organisms must constantly balance biological stability against the energetic costs of over-activating their nervous systems in response to the enormous environmental challenges that living in the wild presents. Thus, regulatory components like BE5.1 might provide three primary points of evolutionary advantages:

Maintaining neural circuits in a permanently heightened state of transcription is highly demanding and energetically expensive. In the wild, networks (such as vesicular transporters, kinases, and transcription factors) are kept in a suppressed state despite the considerable environmental challenges faced. By serving as a genomic “brake”, BE5.1 might prevent cells from prematurely exhausting metabolic resources on unneeded proteins, ensuring that high-level expression is strictly gated.

Unchecked expression of major neurotransmitter machinery poses an inherent physiological risk that might impact on species fitness. In WT hypothalamus, MS triggers a reduction in the expression of genes such as *Slc17a7* (VGLUT1) and *Slc30a3* (ZnT3) thus ensuing an appropriate response that does not affect the health of the animal. However, in KO animals these genes are strongly upregulated which may have detrimental effects as VGLUT1 controls the loading of glutamate into presynaptic vesicles while ZnT3 packages zinc, a critical co-released modulator of post-synaptic NMDA receptors. Without a brake like BE5.1, a glutamatergic surge can lead to localized hyperexcitability, excessive calcium influx via post-synaptic receptors, and metabolic excitotoxicity. BE5.1 might, therefore, protect the structural integrity of the hypothalamus from self-induced, over-excitation tissue damage following environmental challenge thus contributing to resilience.

The hypothalamus must coordinate highly precise neuroendocrine feedback loops. Thus, when BE5.1 element is removed, secondary messenger amplifiers like *Adcy1* and potent neuropeptides like *Cck* and *Cort* are highly expressed. In the wild, exposure to environmental challenge, in the absence of BE5.1, could lead to an unbuffered endocrine loop and an over-reactive hypothalamic-pituitary-adrenal (HPA) axis, causing chronic stress, immune suppression, and reproductive issues. It could therefore be argued that BE5.1 provides an evolutionary cushion against environmental stress that keeps metabolic axes stable during childhood development with impacts on food intake and anxiety pathways.

### Significance for understanding the effects of early life stress in humans

The current study has attempted to model the role of a highly conserved regulatory element, that harbours an obesity and anxiety associated polymorphism, in modulating the effects of early-life stress later in life [21, 22]. Although the data presented is complex and varies between tissues, specific behaviours and sex, the overall picture is that BE5.1 contributes to specific MS induced behaviours by buffering against the overexpression of specific groups of genes thereby maintaining homeostasis and preserving behaviours consistent with survival in a challenging environment. Indeed, one of the ways that allelic variation within BE5.1 may confer susceptibility to the effects of MS is by compromising the binding of specific transcription factors, such as *MEF2c*. Alterations in the ability of MEF2c to bind BE5.1 could compromise the ability of BE5.1 to buffer against the effects of environmental challenge and increasing susceptibility to stress induced neuronal hypersensitivity contributing to susceptibility to anxiety/obesity. Indeed, our preliminary *in-vitro* experiments suggest that there is a significant difference in the response of the A and T alleles of the human BE5.1 to *MEF2c* introduction and PKC stimulation and that the *MEF2c* and *Pkcge* genes are amongst those suppressed in WT animals but upregulated in the hypothalamus of male and female animals. Together, these observations suggest a mechanistic model of how allelic variants of BE5.1 might contribute to weight gain and anxiety in individuals exposed to environmental challenge such as MS via homeostatic autocrine loops.

### Study limitations

Whilst we believe that our study of the role of BE5.1 in modulating the effects of MS on later life gene expression and behaviour reveals critical mechanisms that may have a bearing on human health, the study is not without its limitations. For example, a more thorough analysis of food intake would have complemented the weight gain studies as would determinations of the effects of BE5.1KO and MS on ethanol intake, and these studies are envisaged. Other planned studies include assessments of light/dark cycles on movement and feeding.

Another study limitation is the time difference between exposure to MS, behavioural testing and RNA-seq analysis of gene expression. Thus, changes in the expression of many genes affected by either the KO or MS may have changed in specific tissues by the time that these tissues were collected for analysis. One of the genes most noticeable altered in expression compared to our previous study is BDNF which we previously showed was strongly upregulated in females in response to deletion of BE5.1 and coincided with an increase in anxiety[21]. However, in the current study, whilst we observed a similar increase in anxiety as a result of the KO in females, relatively muted changes in BDNF expression were observed in both males and females. Whilst these observed differences in BDNF expression may be a result of our revised handling procedures, we also recognise that BE5.1 might regulate BDNF expression earlier in development and outside of the timeframe covered by our RNA-seq analysis. We hope to address this limitation by repeating RNA-seq analysis in WT and KO mice just after MS and in adolescent animals.

We also cannot rule out the possibility that, despite being situated in an intron of BDNF, the regulation of BDNF may not be the primary function of BE5.1. This argument is not without precedent as a number of cis-regulatory sequences have been identified within the intronic sequences of genes (*LMBR1* and *FTO*) whose expression is largely independent of these sequences [79, 80]. A further possibility is that BE5.1 deletion may compromise the development of specific regions of the brain thus affecting behaviour and the spectrum of gene expression pertinent to these missing regions. Thus, more study of these lines is required before any direct link between BE5.1 activity and the changes in gene expression observed can be made.

A further limitation of the current study is that, although we provide robust data demonstrating a role for BE5.1 in modulating transcriptome and behavioural response to environmental challenges, and differences in the reaction of the human A and T-alleles of BE5.1 to cellular cues *in-vitro*, we were unable to model the effects of these human alleles on BE5.1 activity *in-vivo.* In the past, addressing this deficit would have been a challenging prospect with the production of humanised mouse models only achievable using embryonic stem-cell targeting, a technically challenging, expensive and time-consuming process. However, with the recent development of adeno-associated virus repair template delivery systems for rapid and targeted modification of mouse zygotes [81] production and analysis of these models has become eminently and rapidly achievable.

### Conclusions

There is mounting evidence that many human conditions arise from variations in the context-dependent CREs within the non-coding genome, that direct the spatial, temporal, and inducible transcription of protein-coding genes essential for normal development and health. Here, we identify a CRE containing a disease-associated SNP that contributes to the neurological homeostasis required for appropriate responses to environmental challenge. Crucially, this work depended on *in-vivo* models, reinforcing the argument that context-dependent CREs cannot be adequately interrogated using *in-vitro* systems [82]. Our previous studies have shown that a substantial subset of these regulatory elements are highly conserved and can be detected through comparative analysis [83], and CRISPR-based genome editing now enables their functional dissection in placental mammalian models. We have also demonstrated that CRE activity can be modified by disease-associated polymorphisms and can be affected by environmental challenge. Although much remains to be understood of the molecular mechanisms underlying BE5.1’s specific role the association between BE5.1 deletion, altered weight gain, and anxiety-related behaviour—together with the accompanying transcriptomic changes—the current study provides a clear path for future investigation into how non-coding regulatory mechanisms shape human health. These findings support continued in-vivo based research into obesity, anxiety, environmental resilience, and the cis-regulatory architecture that governs them.

## Acknowledgements

AMcE, JM, CM and AMcK were funded by a BBSRC project grant (BB/N017544/1) with contributions from Tenovus Scotland Grampian (award G19.08). GH was funded by Medical Research Scotland (PHD-50366-2021) shared by AMcK and EC-D. We thank the staff (Darrin Sheppard, Donna Wallace, Andrew Brown, Scott Mackie, Mark Herbert and Karen Heath) at the Medical Research Facility for their help and excellent advice towards facilitation the completion of these studies.

## Data availability

The datasets generated and/or analysed during the current study are available from the corresponding author on reasonable request.

## Author contributions

**AMcK**; study design, generation of GA animals, analysis, interpretation, writing of manuscript, corresponding author. **AMcE,** study design, animal behavioural analysis, interpretation, writing, first author. **AR, JAJ** and **GR,** study design, animal behavioural studies and manuscript editing. **GH,** DNA cloning, QPCR, luciferase analyses, cell culture. **E C-D, ZH** and **L N van de L,** initial RNA-seq analysis and experimental design. **CH** and **JM,** more extensive RNA-seq analysis, and statistical analysis

## References

1. Green, J. G., McLaughlin, K. A., Berglund, P. A., Gruber, M. J., Sampson, N. A., Zaslavsky, A. M. & Kessler, R. C. (2010) Childhood adversities and adult psychiatric disorders in the national comorbidity survey replication I: associations with first onset of DSM-IV disorders, Arch Gen Psychiatry. 67, 113–23.

2. Felitti, V. J., Anda, R. F., Nordenberg, D., Williamson, D. F., Spitz, A. M., Edwards, V., Koss, M. P. & Marks, J. S. (2019) REPRINT OF: Relationship of Childhood Abuse and Household Dysfunction to Many of the Leading Causes of Death in Adults: The Adverse Childhood Experiences (ACE) Study, Am J Prev Med. 56, 774–786.

3. Nicolaides, N. C., Kanaka-Gantenbein, C. & Pervanidou, P. (2024) Developmental Neuroendocrinology of Early-Life Stress: Impact on Child Development and Behavior, Curr Neuropharmacol. 22, 461–474.

4. Hemmingsson, E. (2014) A new model of the role of psychological and emotional distress in promoting obesity: conceptual review with implications for treatment and prevention, Obes Rev. 15, 769–79.

5. Kaman, A., Devine, J., Erhart, M., Napp, A. K., Reiss, F., Moeller, S., Zoellner, F., Behn, S. & Ravens-Sieberer, U. (2025) Youth Mental Health in Times of Global Crises: Evidence From the German Longitudinal COVID-19 and Psychological Health Study, J Adolesc Health. 77, 1050–1057.

6. Peterson, C., Aslam, M. V., Niolon, P. H., Bacon, S., Bellis, M. A., Mercy, J. A. & Florence, C. (2023) Economic Burden of Health Conditions Associated With Adverse Childhood Experiences Among US Adults, JAMA Netw Open. 6, e2346323.

7. Government, S. (2019) Scottish Health Survey 2019 - volume 1: main report in (Directorate, P. H., ed), Scottish Government,

8. Segenreich, D., Paez, M. S., Regalla, M. A., Fortes, D., Faraone, S. V., Sergeant, J. & Mattos, P. (2015) Multilevel analysis of ADHD, anxiety and depression symptoms aggregation in families, Eur Child Adolesc Psychiatry. 24, 525–36.

9. Kundakovic, M., Lim, S., Gudsnuk, K. & Champagne, F. A. (2013) Sex-specific and strain-dependent effects of early life adversity on behavioral and epigenetic outcomes, Front Psychiatry. 4, 78.

10. Waterhouse, E. G., An, J. J., Orefice, L. L., Baydyuk, M., Liao, G. Y., Zheng, K., Lu, B. & Xu, B. (2012) BDNF promotes differentiation and maturation of adult-born neurons through GABAergic transmission, J Neurosci. 32, 14318–30.

11. Campbell, T. S., Donoghue, K. M., Ghosh, U., Nelson, C. M. & Roth, T. L. (2022) Early Life Stress Affects Bdnf Regulation: A Role for Exercise Interventions, Int J Mol Sci. 23.

12. Aas, M., Dieset, I., Morch, R., Steen, N. E., Hope, S., Reponen, E. J., Laskemoen, J. F., Ueland, T., Aukrust, P., Melle, I., Agartz, I. & Andreassen, O. A. (2019) Reduced brain-derived neurotrophic factor is associated with childhood trauma experiences and number of depressive episodes in severe mental disorders, Schizophrenia Research. 205, 45–50.

13. Sakaue, S., Kanai, M., Tanigawa, Y., Karjalainen, J., Kurki, M., Koshiba, S., Narita, A., Konuma, T., Yamamoto, K., Akiyama, M., Ishigaki, K., Suzuki, A., Suzuki, K., Obara, W., Yamaji, K., Takahashi, K., Asai, S., Takahashi, Y., Suzuki, T., Shinozaki, N., Yamaguchi, H., Minami, S., Murayama, S., Yoshimori, K., Nagayama, S., Obata, D., Higashiyama, M., Masumoto, A., Koretsune, Y., FinnGen, Ito, K., Terao, C., Yamauchi, T., Komuro, I., Kadowaki, T., Tamiya, G., Yamamoto, M., Nakamura, Y., Kubo, M., Murakami, Y., Yamamoto, K., Kamatani, Y., Palotie, A., Rivas, M. A., Daly, M. J., Matsuda, K. & Okada, Y. (2021) A cross-population atlas of genetic associations for 220 human phenotypes, Nat Genet. 53, 1415–1424.

14. Zhu, Z., Guo, Y., Shi, H., Liu, C. L., Panganiban, R. A., Chung, W., O’Connor, L. J., Himes, B. E., Gazal, S., Hasegawa, K., Camargo, C. A., Jr., Qi, L., Moffatt, M. F., Hu, F. B., Lu, Q., Cookson, W. O. C. & Liang, L. (2020) Shared genetic and experimental links between obesity-related traits and asthma subtypes in UK Biobank, J Allergy Clin Immunol. 145, 537–549.

15. Koskeridis, F., Evangelou, E., Said, S., Boyle, J. J., Elliott, P., Dehghan, A. & Tzoulaki, I. (2022) Pleiotropic genetic architecture and novel loci for C-reactive protein levels, Nat Commun. 13, 6939.

16. Wang, H., Zhang, F., Zeng, J., Wu, Y., Kemper, K. E., Xue, A., Zhang, M., Powell, J. E., Goddard, M. E., Wray, N. R., Visscher, P. M., McRae, A. F. & Yang, J. (2019) Genotype-by-environment interactions inferred from genetic effects on phenotypic variability in the UK Biobank, Sci Adv. 5, eaaw3538.

17. Nagel, M., Jansen, P. R., Stringer, S., Watanabe, K., de Leeuw, C. A., Bryois, J., Savage, J. E., Hammerschlag, A. R., Skene, N. G., Munoz-Manchado, A. B., and Me Research, T., White, T., Tiemeier, H., Linnarsson, S., Hjerling-Leffler, J., Polderman, T. J. C., Sullivan, P. F., van der Sluis, S. & Posthuma, D. (2018) Meta-analysis of genome-wide association studies for neuroticism in 449,484 individuals identifies novel genetic loci and pathways, Nat Genet. 50, 920–927.

18. Nagel, M., Watanabe, K., Stringer, S., Posthuma, D. & van der Sluis, S. (2018) Item-level analyses reveal genetic heterogeneity in neuroticism, Nat Commun. 9, 905.

19. Hill, W. D., Weiss, A., Liewald, D. C., Davies, G., Porteous, D. J., Hayward, C., McIntosh, A. M., Gale, C. R. & Deary, I. J. (2020) Genetic contributions to two special factors of neuroticism are associated with affluence, higher intelligence, better health, and longer life, Mol Psychiatry. 25, 3034–3052.

20. Hing, B., Davidson, S., Lear, M., Breen, G., Quinn, J., McGuffin, P. & MacKenzie, A. (2012) A polymorphism associated with depressive disorders differentially regulates brain derived neurotrophic factor promoter IV activity, Biol Psychiatry. 71, 618–26.

21. McEwan, A. R., Hing, B., Erickson, J. C., Hutchings, G., Urama, C., Norton-Hughes, E., D’Ippolito, M., Berry, S., Delibegovic, M., Grassmann, F. & MacKenzie, A. (2024) An ancient polymorphic regulatory region within the BDNF gene associated with obesity modulates anxiety-like behaviour in mice and humans, Mol Psychiatry.

22. Speliotes, E. K. Willer, C. J. Berndt, S. I. Monda, K. L. Thorleifsson, G. Jackson, A. U. Lango Allen, H. Lindgren, C. M. Luan, J. Magi, R. Randall, J. C. Vedantam, S. Winkler, T. W. Qi, L. Workalemahu, T. Heid, I. M. Steinthorsdottir, V. Stringham, H. M. Weedon, M. N. Wheeler, E. Wood, A. R. Ferreira, T. Weyant, R. J. Segre, A. V. Estrada, K. Liang, L. Nemesh, J. Park, J. H. Gustafsson, S. Kilpelainen, T. O. Yang, J. Bouatia-Naji, N. Esko, T. Feitosa, M. F. Kutalik, Z. Mangino, M. Raychaudhuri, S. Scherag, A. Smith, A. V. Welch, R. Zhao, J. H. Aben, K. K. Absher, D. M. Amin, N. Dixon, A. L. Fisher, E. Glazer, N. L. Goddard, M. E. Heard-Costa, N. L. Hoesel, V. Hottenga, J. J. Johansson, A. Johnson, T. Ketkar, S. Lamina, C. Li, S. Moffatt, M. F. Myers, R. H. Narisu, N. Perry, J. R. Peters, M. J. Preuss, M. Ripatti, S. Rivadeneira, F. Sandholt, C. Scott, L. J. Timpson, N. J. Tyrer, J. P. van Wingerden, S. Watanabe, R. M. White, C. C. Wiklund, F. Barlassina, C. Chasman, D. I. Cooper, M. N. Jansson, J. O. Lawrence, R. W. Pellikka, N. Prokopenko, I. Shi, J. Thiering, E. Alavere, H. Alibrandi, M. T. Almgren, P. Arnold, A. M. Aspelund, T. Atwood, L. D. Balkau, B. Balmforth, A. J. Bennett, A. J. Ben-Shlomo, Y. Bergman, R. N. Bergmann, S. Biebermann, H. Blakemore, A. I. Boes, T. Bonnycastle, L. L. Bornstein, S. R. Brown, M. J. Buchanan, T. A. et al. (2010) Association analyses of 249,796 individuals reveal 18 new loci associated with body mass index, Nat Genet. 42, 937–48.

23. Gouveia, K. & Hurst, J. L. (2019) Improving the practicality of using non-aversive handling methods to reduce background stress and anxiety in laboratory mice, Sci Rep. 9, 20305.

24. Szabo, J., Renczes, E., Borbelyova, V., Ostatnikova, D. & Celec, P. (2024) Assessing sociability using the Three-Chamber Social Interaction Test and the Reciprocal Interaction Test in a genetic mouse model of ASD, Behav Brain Funct. 20, 24.

25. Kent, W. J., Sugnet, C. W., Furey, T. S., Roskin, K. M., Pringle, T. H., Zahler, A. M. & Haussler, D. (2002) The human genome browser at UCSC, Genome Res. 12, 996–1006.

26. Navarro Gonzalez, J., Zweig, A. S., Speir, M. L., Schmelter, D., Rosenbloom, K. R., Raney, B. J., Powell, C. C., Nassar, L. R., Maulding, N. D., Lee, C. M., Lee, B. T., Hinrichs, A. S., Fyfe, A. C., Fernandes, J. D., Diekhans, M., Clawson, H., Casper, J., Benet-Pages, A., Barber, G. P., Haussler, D., Kuhn, R. M., Haeussler, M. & Kent, W. J. (2021) The UCSC Genome Browser database: 2021 update, Nucleic Acids Res. 49, D1046–D1057.

27. Li, H. (2018) Minimap2: pairwise alignment for nucleotide sequences, Bioinformatics. 34, 3094–3100.

28. Liao, Y., Smyth, G. K. & Shi, W. (2014) featureCounts: an efficient general purpose program for assigning sequence reads to genomic features, Bioinformatics. 30, 923–30.

29. O’Leary, N. A., Wright, M. W., Brister, J. R., Ciufo, S., Haddad, D., McVeigh, R., Rajput, B., Robbertse, B., Smith-White, B., Ako-Adjei, D., Astashyn, A., Badretdin, A., Bao, Y., Blinkova, O., Brover, V., Chetvernin, V., Choi, J., Cox, E., Ermolaeva, O., Farrell, C. M., Goldfarb, T., Gupta, T., Haft, D., Hatcher, E., Hlavina, W., Joardar, V. S., Kodali, V. K., Li, W., Maglott, D., Masterson, P., McGarvey, K. M., Murphy, M. R., O’Neill, K., Pujar, S., Rangwala, S. H., Rausch, D., Riddick, L. D., Schoch, C., Shkeda, A., Storz, S. S., Sun, H., Thibaud-Nissen, F., Tolstoy, I., Tully, R. E., Vatsan, A. R., Wallin, C., Webb, D., Wu, W., Landrum, M. J., Kimchi, A., Tatusova, T., DiCuccio, M., Kitts, P., Murphy, T. D. & Pruitt, K. D. (2016) Reference sequence (RefSeq) database at NCBI: current status, taxonomic expansion, and functional annotation, Nucleic Acids Res. 44, D733–45.

30. Torres-Berrio, A., Bortolami, A., Pena, C. J. & Nestler, E. J. (2025) Neurobiology of resilience to early life stress, Neuropsychopharmacology.

31. Own, L. S., Iqbal, R. & Patel, P. D. (2013) Maternal separation alters serotonergic and HPA axis gene expression independent of separation duration in mice, Brain Res. 1515, 29–38.

32. George, E. D., Bordner, K. A., Elwafi, H. M. & Simen, A. A. (2010) Maternal separation with early weaning: a novel mouse model of early life neglect, BMC Neurosci. 11, 123.

33. Tao, E., Long, G., Yang, T., Chen, B., Guo, R., Ye, D., Fang, M. & Jiang, M. (2021) Maternal Separation Induced Visceral Hypersensitivity Evaluated via Novel and Small Size Distention Balloon in Post-weaning Mice, Front Neurosci. 15, 803957.

34. Baugher, B. J. & Sachs, B. D. (2022) Early life maternal separation induces sex-specific antidepressant-like responses but has minimal effects on adult stress susceptibility in mice, Front Behav Neurosci. 16, 941884.

35. Angoa-Perez, M., Kane, M. J., Briggs, D. I., Francescutti, D. M. & Kuhn, D. M. (2013) Marble burying and nestlet shredding as tests of repetitive, compulsive-like behaviors in mice, J Vis Exp, 50978.

36. Thomas, A., Burant, A., Bui, N., Graham, D., Yuva-Paylor, L. A. & Paylor, R. (2009) Marble burying reflects a repetitive and perseverative behavior more than novelty-induced anxiety, Psychopharmacology (Berl). 204, 361–73.

37. Porcher, C., Medina, I. & Gaiarsa, J. L. (2018) Mechanism of BDNF Modulation in GABAergic Synaptic Transmission in Healthy and Disease Brains, Front Cell Neurosci. 12, 273.

38. Amrutkar, M., Cansby, E., Chursa, U., Nunez-Duran, E., Chanclon, B., Stahlman, M., Friden, V., Manneras-Holm, L., Wickman, A., Smith, U., Backhed, F., Boren, J., Howell, B. W. & Mahlapuu, M. (2015) Genetic Disruption of Protein Kinase STK25 Ameliorates Metabolic Defects in a Diet-Induced Type 2 Diabetes Model, Diabetes. 64, 2791–804.

39. Chelban, V., Alsagob, M., Kloth, K., Chirita-Emandi, A., Vandrovcova, J., Maroofian, R., Davagnanam, I., Bakhtiari, S., AlSayed, M. D., Rahbeeni, Z., AlZaidan, H., Malintan, N. T., Johannsen, J., Efthymiou, S., Ghayoor Karimiani, E., Mankad, K., Al-Shahrani, S. A., Beiraghi Toosi, M., AlShammari, M., Groppa, S., Haridy, N. A., AlQuait, L., Qari, A., Huma, R., Salih, M. A., Almass, R., Almutairi, F. B., Hamad, M. H., Alorainy, I. A., Ramzan, K., Imtiaz, F., Puiu, M., Kruer, M. C., Bierhals, T., Wood, N. W., Colak, D., Houlden, H. & Kaya, N. (2020) Genetic and phenotypic characterization of NKX6-2-related spastic ataxia and hypomyelination, Eur J Neurol. 27, 334–342.

40. Walker, L. C., Hand, L. J., Letherby, B., Huckstep, K. L., Campbell, E. J. & Lawrence, A. J. (2021) Cocaine and amphetamine regulated transcript (CART) signalling in the central nucleus of the amygdala modulates stress-induced alcohol seeking, Neuropsychopharmacology. 46, 325–333.

41. Bowers, B. J., Collins, A. C., Tritto, T. & Wehner, J. M. (2000) Mice lacking PKC gamma exhibit decreased anxiety, Behav Genet. 30, 111–21.

42. Kansra, A. R., Lakkunarajah, S. & Jay, M. S. (2020) Childhood and Adolescent Obesity: A Review, Front Pediatr. 8, 581461.

43. Bartsch, V. B., Lord, J. S., Diering, G. H. & Zylka, M. J. (2020) Mania- and anxiety-like behavior and impaired maternal care in female diacylglycerol kinase eta and iota double knockout mice, Genes Brain Behav. 19, e12570.

44. Davis, M. M., Olausson, P., Greengard, P., Taylor, J. R. & Nairn, A. C. (2012) Regulator of calmodulin signaling knockout mice display anxiety-like behavior and motivational deficits, Eur J Neurosci. 35, 300–8.

45. Bourin, M. (1998) Cholecystokinin as a target for neuropsychiatric drugs, Drug News Perspect. 11, 342–9.

46. Zhang, R., Asai, M., Mahoney, C. E., Joachim, M., Shen, Y., Gunner, G. & Majzoub, J. A. (2017) Loss of hypothalamic corticotropin-releasing hormone markedly reduces anxiety behaviors in mice, Mol Psychiatry. 22, 733–744.

47. Gray, J. M., Vecchiarelli, H. A., Morena, M., Lee, T. T., Hermanson, D. J., Kim, A. B., McLaughlin, R. J., Hassan, K. I., Kuhne, C., Wotjak, C. T., Deussing, J. M., Patel, S. & Hill, M. N. (2015) Corticotropin-releasing hormone drives anandamide hydrolysis in the amygdala to promote anxiety, J Neurosci. 35, 3879–92.

48. Miao, X., Fu, Z., Luo, X., Wang, J., Yuan, L., Zhao, S., Feng, Y., Huang, S. & Xiao, S. (2024) A study on the correlations of PRL levels with anxiety, depression, sleep, and self-efficacy in patients with prolactinoma, Front Endocrinol (Lausanne). 15, 1369729.

49. Ghazi Ghanim, K., Saab Kadhim, M., Hameed Abed Ali, B. & Jawad, R. A. (2023) The Relation between Increasing Anxiety and Prolactin-Releasing Peptide in Rats, Arch Razi Inst. 78, 181–184.

50. Jaszberenyi, M., Thurzo, B., Bagosi, Z., Vecsei, L. & Tanaka, M. (2024) The Orexin/Hypocretin System, the Peptidergic Regulator of Vigilance, Orchestrates Adaptation to Stress, Biomedicines. 12.

51. Valentinuzzi, V. S., Kolker, D. E., Vitaterna, M. H., Shimomura, K., Whiteley, A., Low-Zeddies, S., Turek, F. W., Ferrari, E. A., Paylor, R. & Takahashi, J. S. (1998) Automated measurement of mouse freezing behavior and its use for quantitative trait locus analysis of contextual fear conditioning in (BALB/cJ x C57BL/6J)F2 mice, Learn Mem. 5, 391–403.

52. Handa, R. J., Sheng, J. A., Castellanos, E. A., Templeton, H. N. & McGivern, R. F. (2022) Sex Differences in Acute Neuroendocrine Responses to Stressors in Rodents and Humans, Cold Spring Harb Perspect Biol. 14.

53. Kember, R. L., Dempster, E. L., Lee, T. H., Schalkwyk, L. C., Mill, J. & Fernandes, C. (2012) Maternal separation is associated with strain-specific responses to stress and epigenetic alterations to Nr3c1, Avp, and Nr4a1 in mouse, Brain Behav. 2, 455–67.

54. Romeo, R. D., Mueller, A., Sisti, H. M., Ogawa, S., McEwen, B. S. & Brake, W. G. (2003) Anxiety and fear behaviors in adult male and female C57BL/6 mice are modulated by maternal separation, Horm Behav. 43, 561–7.

55. Fan, W., Ellacott, K. L., Halatchev, I. G., Takahashi, K., Yu, P. & Cone, R. D. (2004) Cholecystokinin-mediated suppression of feeding involves the brainstem melanocortin system, Nat Neurosci. 7, 335–6.

56. Tractenberg, S. G., Levandowski, M. L., de Azeredo, L. A., Orso, R., Roithmann, L. G., Hoffmann, E. S., Brenhouse, H. & Grassi-Oliveira, R. (2016) An overview of maternal separation effects on behavioural outcomes in mice: Evidence from a four-stage methodological systematic review, Neurosci Biobehav Rev. 68, 489–503.

57. Murgatroyd, C., Patchev, A. V., Wu, Y., Micale, V., Bockmuhl, Y., Fischer, D., Holsboer, F., Wotjak, C. T., Almeida, O. F. & Spengler, D. (2009) Dynamic DNA methylation programs persistent adverse effects of early-life stress, Nat Neurosci. 12, 1559–66.

58. Own, L. S. & Patel, P. D. (2013) Maternal behavior and offspring resiliency to maternal separation in C57Bl/6 mice, Horm Behav. 63, 411–7.

59. Zhang, H., Xiao, X., Pan, Z. & Dokudovskaya, S. (2025) mTOR signaling networks: mechanistic insights and translational frontiers in disease therapeutics, Signal Transduct Target Ther. 10, 428.

60. Meister, B. (2007) Neurotransmitters in key neurons of the hypothalamus that regulate feeding behavior and body weight, Physiol Behav. 92, 263–71.

61. McAllister, B. B., Wright, D. K., Wortman, R. C., Shultz, S. R. & Dyck, R. H. (2018) Elimination of vesicular zinc alters the behavioural and neuroanatomical effects of social defeat stress in mice, Neurobiol Stress. 9, 199–213.

62. Momin, A. A., Mendes, T., Barthe, P., Faure, C., Hong, S., Yu, P., Kadare, G., Jaremko, M., Girault, J. A., Jaremko, L. & Arold, S. T. (2022) PYK2 senses calcium through a disordered dimerization and calmodulin-binding element, Commun Biol. 5, 800.

63. de Pins, B., Mendes, T., Giralt, A. & Girault, J. A. (2021) The Non-receptor Tyrosine Kinase Pyk2 in Brain Function and Neurological and Psychiatric Diseases, Front Synaptic Neurosci. 13, 749001.

64. Del Turco, D., Paul, M. H., Rietsche, M., Schlaudraff, J., Kreuzer, A., Mittag, M., Drakew, A., Jedlicka, P., Bas-Orth, C., Roeper, J. & Deller, T. (2026) Overexpression of Synaptopodin increases the number of spine apparatuses and active synapses of dentate granule cells, Sci Rep. 16.

65. Ormond, C., Cap, M., Chang, Y. C., Ryan, N., Chavira, D., Williams, K., Grant, J. E., Mathews, C., Heron, E. A. & Corvin, A. (2026) Rare protein-disrupting variants in NPY5R, DLGAP1 and MAPK8IP3 segregate with OCD in two multiplex pedigrees, medRxiv.

66. Minagawa, K., Hayakawa, T., Akimoto, H., Nagashima, T., Takahashi, Y. & Asai, S. (2025) Late development of OCD-like phenotypes in Dlgap1 knockout mice, Psychopharmacology (Berl). 242, 215–231.

67. Yang, M., Ding, Q., Zhang, M., Moon, C. & Wang, H. (2020) Forebrain overexpression of type 1 adenylyl cyclase promotes molecular stability and behavioral resilience to physical stress, Neurobiol Stress. 13, 100237.

68. Bondar, N. P., Lepeshko, A. A. & Reshetnikov, V. V. (2018) Effects of Early-Life Stress on Social and Anxiety-Like Behaviors in Adult Mice: Sex-Specific Effects, Behav Neurol. 2018, 1538931.

69. Endo, N., Makinodan, M., Mannari-Sasagawa, T., Horii-Hayashi, N., Somayama, N., Komori, T., Kishimoto, T. & Nishi, M. (2021) The effects of maternal separation on behaviours under social-housing environments in adult male C57BL/6 mice, Sci Rep. 11, 527.

70. Millstein, R. A. & Holmes, A. (2007) Effects of repeated maternal separation on anxiety- and depression-related phenotypes in different mouse strains, Neurosci Biobehav Rev. 31, 3–17.

71. Destree, L., Brierley, M. E., Albertella, L., Jobson, L. & Fontenelle, L. F. (2021) The effect of childhood trauma on the severity of obsessive-compulsive symptoms: A systematic review, J Psychiatr Res. 142, 345–360.

72. Han, E., Canada, K. A., Puglia, M. H., Pelphrey, K. A. & Evans, T. M. (2025) The Convergence of Early-Life Stress and Autism Spectrum Disorder on the Epigenetics of Genes Key to the HPA Axis, Biology (Basel). 15.

73. Hemmings, S. M., Lochner, C., van der Merwe, L., Cath, D. C., Seedat, S. & Stein, D. J. (2013) BDNF Val66Met modifies the risk of childhood trauma on obsessive-compulsive disorder, J Psychiatr Res. 47, 1857–63.

74. Li, P., Cheng, J., Gu, Q., Wang, P., Lin, Z., Fan, Q., Chen, J. & Wang, Z. (2021) Intermediation of perceived stress between early trauma and plasma M/P ratio levels in obsessive-compulsive disorder patients, J Affect Disord. 285, 105–111.

75. Hu, P., Maita, I., Phan, M. L., Gu, E., Kwok, C., Dieterich, A., Gergues, M. M., Yohn, C. N., Wang, Y., Zhou, J. N., Qi, X. R., Swaab, D. F., Pang, Z. P., Lucassen, P. J., Roepke, T. A. & Samuels, B. A. (2020) Early-life stress alters affective behaviors in adult mice through persistent activation of CRH-BDNF signaling in the oval bed nucleus of the stria terminalis, Transl Psychiatry. 10, 396.

76. de Brouwer, G., Fick, A., Harvey, B. H. & Wolmarans, W. (2019) A critical inquiry into marble-burying as a preclinical screening paradigm of relevance for anxiety and obsessive-compulsive disorder: Mapping the way forward, Cogn Affect Behav Neurosci. 19, 1–39.

77. Abraham, M., Schmerder, K., Hedtstuck, M., Bosing, K., Mundorf, A. & Freund, N. (2023) Maternal separation and its developmental consequences on anxiety and parvalbumin interneurons in the amygdala, J Neural Transm (Vienna). 130, 1167–1175.

78. Taylor, G. T., Lerch, S. & Chourbaji, S. (2017) Marble burying as compulsive behaviors in male and female mice, Acta Neurobiol Exp (Wars). 77, 254–260.

79. Smemo, S., Tena, J. J., Kim, K. H., Gamazon, E. R., Sakabe, N. J., Gomez-Marin, C., Aneas, I., Credidio, F. L., Sobreira, D. R., Wasserman, N. F., Lee, J. H., Puviindran, V., Tam, D., Shen, M., Son, J. E., Vakili, N. A., Sung, H. K., Naranjo, S., Acemel, R. D., Manzanares, M., Nagy, A., Cox, N. J., Hui, C. C., Gomez-Skarmeta, J. L. & Nobrega, M. A. (2014) Obesity-associated variants within FTO form long-range functional connections with IRX3, Nature. 507, 371–5.

80. Lettice, L. A., Horikoshi, T., Heaney, S. J., van Baren, M. J., van der Linde, H. C., Breedveld, G. J., Joosse, M., Akarsu, N., Oostra, B. A., Endo, N., Shibata, M., Suzuki, M., Takahashi, E., Shinka, T., Nakahori, Y., Ayusawa, D., Nakabayashi, K., Scherer, S. W., Heutink, P., Hill, R. E. & Noji, S. (2002) Disruption of a long-range cis-acting regulator for Shh causes preaxial polydactyly, Proc Natl Acad Sci U S A. 99, 7548–53.

81. Moncaut, N. (2024) Streamlining mouse genome editing by integrating AAV repair template delivery and CRISPR-Cas electroporation, Lab Anim (NY). 53, 115–116.

82. MacKenzie, A., Hay, E. A. & McEwan, A. R. (2022) Context-dependant enhancers as a reservoir of functional polymorphisms and epigenetic markers linked to alcohol use disorders and comorbidities, Addict Neurosci. 2, None.

83. McEwan, A., Rattray, A., Hutchings, G., Hay, E., Murgatroyd, C. & MacKenzie, A. (2025) Exploring the roles of conserved context-dependent cis-regulatory elements (cdCREs) in multicellularity, human health and disease, FEBS J. 292, 6163–6181.

